# Parallel evolution between genomic segments of seasonal human influenza viruses reveals RNA-RNA relationships

**DOI:** 10.1101/2021.01.14.426718

**Authors:** Jennifer E. Jones, Valerie Le Sage, Gabriella H. Padovani, Michael Calderon, Erik S. Wright, Seema S. Lakdawala

## Abstract

The influenza A virus (IAV) genome consists of eight negative-sense viral RNA (vRNA) segments that are selectively assembled into progeny virus particles through RNA-RNA interactions. To identify relationships between vRNA segments, we examined parallel evolution between vRNA segments of seasonal human IAV, finding that evolutionary relationships between vRNA segments differ between subtypes and antigenically-shifted strains. Intersegmental relationships were distinct between H3N2 and H1N1 viruses, but largely conserved over time in H3N2 viruses. However, parallel evolution of vRNA segments diverged between H1N1 strains isolated before and after the 2009 pandemic. Surprisingly, intersegmental relationships were not driven solely by protein sequence, which is potentially indicative of RNA-RNA driven coevolution. Colocalization of highly coevolved vRNA segments was enriched over other pairs at the nuclear periphery during a productive viral infection. This study illustrates how phylogenetics can be applied to interrogate putative RNA interactions underlying selective assembly of IAV.

## Introduction

Genetic variation is ubiquitious in RNA viruses. The rapid evolution underlying this variation can occur as a result of mutation, recombination, or reassortment, with major consequences for human disease (Andino & Domingo, 2015). In the case of influenza virus, these consequences include poor vaccine efficacy rates, immune escape, antiviral resistance, and the emergence of novel strains (Lyons & Lauring, 2018). Within the past century, influenza A virus (IAV) pandemics occurred in 1918 (H1N1), 1957 (H2N2), 1968 (H3N2), and 2009 (H1N1) (Neumann, Noda, & Kawaoka, 2009; Paules & Subbarao, 2017; Short, Kedzierska, & van de Sandt, 2018). Each of the last three influenza pandemics have been attributed to a reassortant strain composed of a novel combination of the eight viral RNA (vRNA) segments of the influenza virus genome (Neumann et al., 2009). Thus, the emergence of pandemic strains is marked by a concommittent alteration in the influenza virus genome triggered by new genetic diversity.

Public health measures to limit the impact of influenza virus outbreaks would benefit from the ability to predict reassortment between circulating influenza viruses. Genetic mutation is driven by stochastic processes and is therefore difficult to predict (Andino & Domingo, 2015). In contrast, reassortment is restricted by a number of factors. As reassortment must occur between two strains coinfecting the same cell, the spatiotemporal dynamics of coinfection as well as compatibility of RNA packaging signals can impede reassortment (Lowen, 2017; Richard, Herfst, Tao, Jacobs, & Lowen, 2018). One method for identifying potential reassortant viruses is to examine intracellular assembly of vRNA segments through biochemical techniques. Intersegmental RNA-RNA interactions have been proposed to facilitate selective assembly and packaging of eight unique vRNA segments into a progeny virion and could pose a significant hurdle to reassortment (Gavazzi et al., 2013; Noda et al., 2006). It is consequently imperative to identify the evolutionary constraints imposed by intersegmental vRNA interactions, as this may help predict future influenza pandemics.

Epistasis within and between genes imposes evolutionary constraints that can be shaped by a number of factors, including the function, stability, or interactions between individual RNA or protein (Sardi & Gasch, 2018). Probabilistic models have revealed several destabilizing mutations in the influenza virus nucleoprotein (NP) that became fixed as a result of counterbalancing epistasis that improves NP protein stability (Gong, Suchard, & Bloom, 2013). These destabilizing mutations occur within T cell epitopes of NP that may be important for immune escape (Gong et al., 2013). Stabilizing epistasis was similarly instrumental to the emergence of oseltamivir-resistance mutations in the influenza neuraminidase (NA) (Bloom, Gong, & Baltimore, 2010). The rise of oseltamivir-resistance mutations in NA spurred investigation of parallel evolution between NA and hemagglutinin (HA), demonstrating that mutations in HA may have facilitated acquisition of oseltamivir-resistance mutations in NA (Jang & Bae, 2018; Kryazhimskiy, Dushoff, Bazykin, & Plotkin, 2011; Neverov, Kryazhimskiy, Plotkin, & Bazykin, 2015). These approaches have great potential, yet the current focus surrounds constraints on protein interactions (Escalera-Zamudio et al., 2020). Such methodologies could be further employed to investigate epistasis arising from RNA-RNA interactions. Mounting evidence from our group and others suggests that direct intermolecular interactions between vRNA segments coordinate selective assembly (Dadonaite et al., 2019; Le Sage et al., 2020). Genomic assembly contributes to heterogeneity in progeny viruses and could determine the fitness of reassortant strains after coinfection (Brooke, 2017; Lowen, 2017). Therefore, parallel evolution between vRNA segments arising from RNA interactions could reveal epistatic constraints on genetic reassortment.

In this study, we set out to combine phylogenetics and molecular biology to examine parallel evolution across vRNA segments genome-wide in seasonal human influenza viruses and identify potential epistatic relationships between vRNA segments. Unlike previous studies, our objective was to identify vRNA segments that might play key roles in genomic assembly. To evaluate phylogenetic relationships among vRNA segments we relied upon the Robinson-Foulds distance (*d*), a measure of topological distance between trees (Robinson, 1981). This method determines the number of branch partitions that are not shared between two trees (Robinson, 1981) and is therefore a quantitative measure of the topological distance between phylogenies. Higher values of *d* correspond to greater topological distance, with a *d* value of 0 indicating that two trees are topologically equivalent. We hypothesized that *d* would vary in accordance with the degree of parallel evolution between genome segments, and sought to determine whether any observed parallel evolution arose from RNA interactions.

## Results

### Phylogenetic relationships between vRNA segments are not uniform in H3N2 viruses

Influenza A virus H1N1 and H3N2 subtypes have cocirculated in human populations since 1977 (Neumann et al., 2009). In order to explore vRNA segment relationships in seasonal human IAV strains over time, we examined parallel evolution between vRNA segments in viruses representative of each subtype. We used four sets of seasonal IAV strains: two sets of H3N2 viruses from 1995-2004 and 2005-2014, and two sets of H1N1 viruses from 2000-2008 and 2010-2018 (**Table 1**). Bracketing the H3N2 viruses into two time intervals permitted investigation of the conservation of vRNA relationships over time in antigenically drifted H3N2 viruses. We took a similar approach with H1N1 viruses, bracketing instead on the antigenic shift event in 2009 with the emergence of the pandemic swine-origin H1N1 virus. Comparison of vRNA relationships in pre-pandemic (2000-2008) and post-pandemic (2010-2018) H1N1 viruses could reveal vRNA relationships from viruses of two distinct lineages or, alternatively, uncover vRNA relationships that remain conserved despite the potential for swapping of vRNA segments across multiple host species.

**Table 1.** Influenza A virus strain datasets. Human H1N1 or H3N2 virus sequences for which full-length sequences are available (Influenza Research Database). Representative sequences were selected for further analysis by clustering. ‘Final Clusters’ indicates the number of clusters after small clusters were collapsed or omitted.

| Subtype | Time Period | Total Strains | Clusters with >97% identity | Final Clusters |
| --- | --- | --- | --- | --- |
| H3N2 | 1995-2004 | 1,026 | 16 | 12 |
|  | 2005-2014 | 3,879 | 17 | 12 |
| H1N1 | 2000-2008 | 821 | 11 | 9 |
|  | 2010-2018 | 4,072 | 14 | 9 |

Our approach, outlined in **Figure 1**, examines evolutionary relationships between vRNA segments. We began our investigation with all H3N2 viruses for which full-length sequence information was available in the Influenza Research Database, yielding 1,026 H3N2 viruses from 1995-2004 and 3,879 H3N2 viruses from 2005-2014 (**Table 1**). However, reconstructing phylogenetic trees from all available sequences was disadvantageous, as a preliminary analysis of three-hundred sequences suggested that a great deal of phylogenetic variation could not be statistically supported by bootstrapping (branch support < 70). Instead, we used a clustering approach to select representative strains that would produce more statistically robust trees. We first concatenated sequences from all strains into full-length genomes from which we built alignments (**Figure 1A**) and clustered into operational taxonomic units on a neighbor-joining species tree (**Figure 1B**). Despite the fact that fewer full-length influenza virus genomic sequences were available prior to the 2000s, our approach resulted in a similar number of clusters within a subtype (**Table 1**), consistent with the notion that increased sequencing has led to more closely related sequences in public databases.

**Figure 1.**
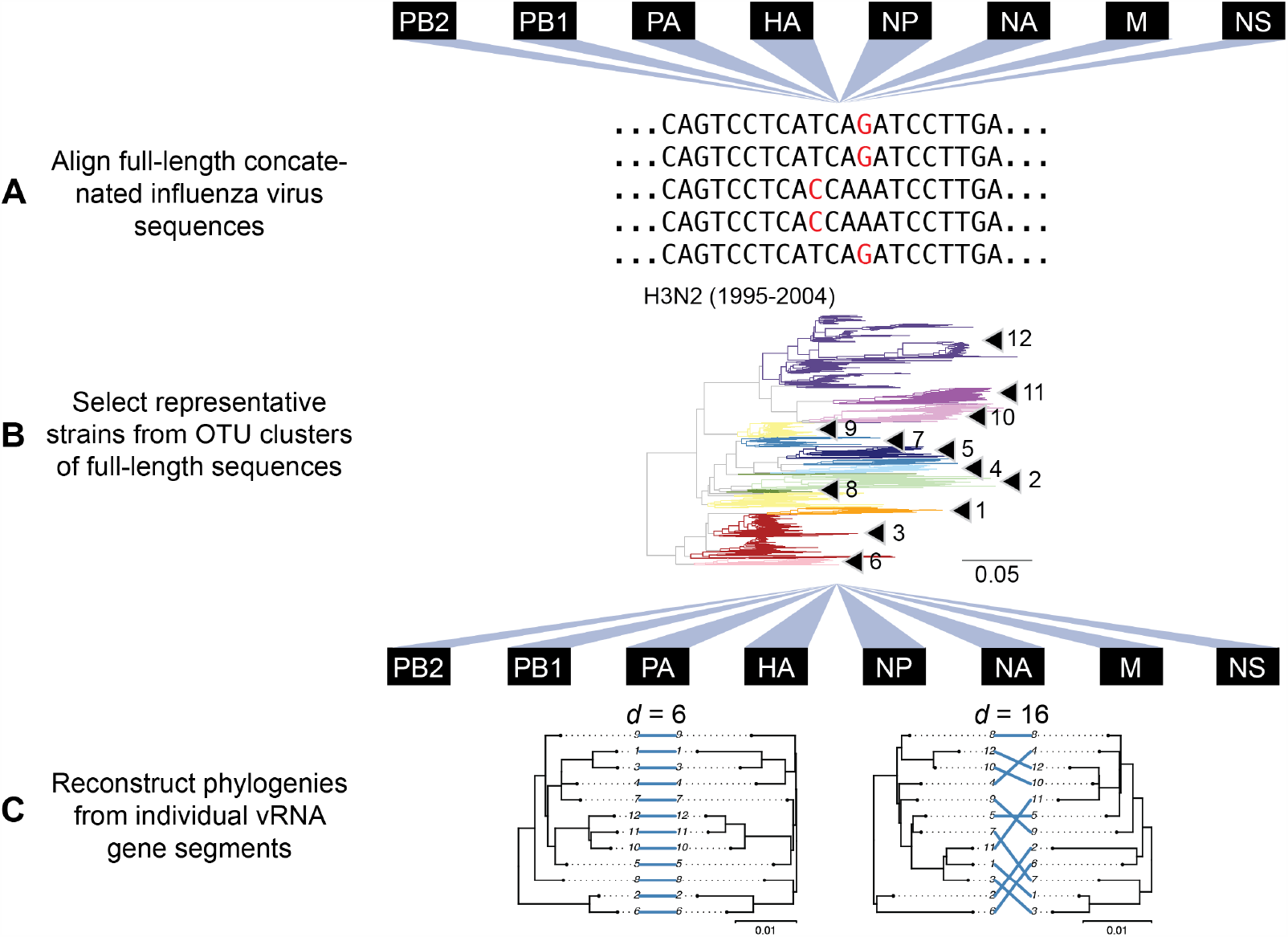
Experimental overview. **A**, Human H3N2 or H1N1 virus sequences were downloaded from the Influenza Research Database and subset into two time periods each: 1995-2004 and 2005-2014 (H3N2 viruses); 2000-2008 and 2010-2018 (H1N1 viruses). The H3N2 virus dataset (1995-2004) is illustrated here. All eight vRNA segments from each strain were concatenated into a full-length genome from which alignments were made. B, A species tree was built grouping strains into operational taxonomic units (OTU clusters) with at least 97% sequence identity. Arrowheads denote clusters 1-12. Seven replicate strains were randomly selected from these clusters for further analysis. C, Full-length genomic sequences were partitioned into individual gene sequence alignments and maximum-likelihood phylogenetic trees were reconstructed from each vRNA gene segment in each replicate. The Robinson-Foulds distances (d) and tanglegrams were examined from each pair of phylogenies. Left, a pair of highly congruent trees with a low *d* value. Right, a pair of discordant trees with a high *d* value. Scale bars indicate percent divergence.

The primary objective behind clustering was to reduce variation between trees that was not statistically supported by bootstrapping. The cutoff for sequence identity during clustering of the species tree was therefore an important consideration because it controlled how much variation remained in our trees. As expected, higher cutoffs (98-99% sequence identity) yielded species trees with more clusters containing fewer members while lower cutoffs (95-96% sequence identity) contained increasingly fewer clusters with more members grouped in each cluster. We selected a cutoff of 97% sequence identity based on the observation that it produced vRNA trees with an intermediate number of clusters (16-17 clusters in each species tree) but enough members in each cluster to provide multiple representative strains for comparison. This process resulted in seven replicate trees for each of the eight vRNA segments, for a total of 56 trees analyzed from each set of H3N2 viruses (**Figure 1C** and **Supplemental Figure 1**).

Concordance between phylogenetic trees built from different RNA segments is expected to be highest when there are strong positive epistatic interactions between encoded protein or RNA complexes (Kryazhimskiy et al., 2011; Neverov et al., 2015). The PB1 and PA proteins are subunits of the heterotrimeric polymerase complex and would be expected to exhibit positive epistasis (Fodor, 2013), whereas PB1 and HA do not share any known protein function. Trees built from one replicate of the PB1 and PA phylogenies from H3N2 (2005-2014) had a low *d* value of 6, suggesting parallel evolution occurs between these two genes (**Figure 2A**). By comparison, the PB1 and HA phylogenies from the same replicate had a *d* value of 14 (**Figure 2B**), suggesting that parallel evolution between PB1 and PA is stronger than between PB1 and HA. Thus, tree concordance recapitulates anticipated protein-driven parallel evolution between two influenza proteins.

**Figure 2.**
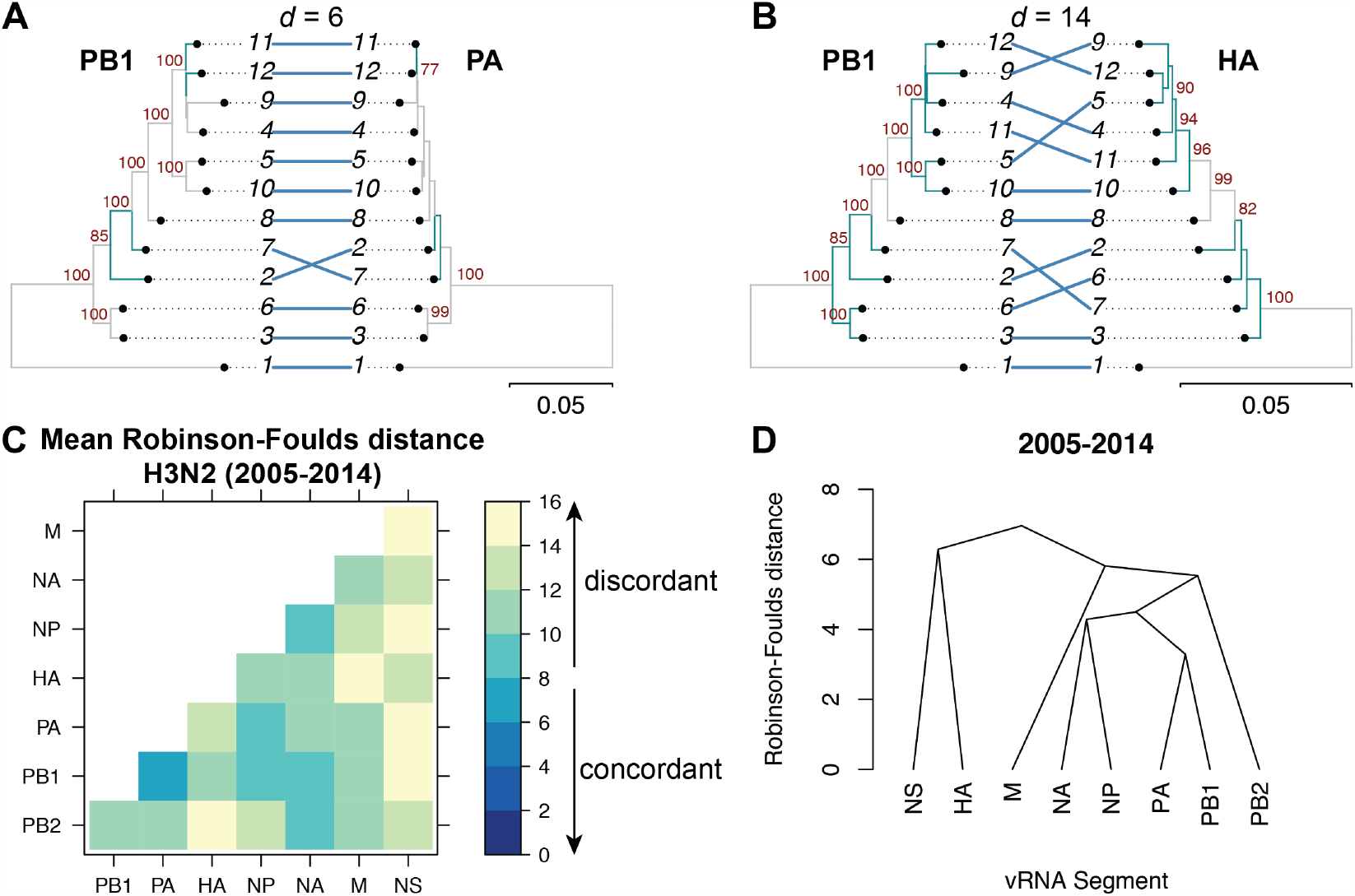
Coevolution between vRNA segments is hierarchical in H3N2 viruses from 2005-2014. Seven replicate maximum-likelihood phylogenies were reconstructed for each vRNA gene segment from human H3N2 virus sequences (2005-2014) as described in Figure 1. **A-B**, Tanglegrams representative of concordant (PB1 and PA gene segments) **(A)** or discordant (PB1 and HA gene segments) phylogenies **(B)** are shown from replicate 1 with discordant branches highlighted in turquoise. Robinson-Foulds distances (*d*) are shown above the tanglegram. Intersecting lines map leaves on the left tree to the corresponding leaves on the right. Strains are coded by cluster number; strain identities can be found in Supplemental Table 2. Bootstrap values greater than 70 are shown in red. Scale bars indicate percent divergence. **C**, Pairwise d values were calculated between each pair of phylogenies in each replicate and the mean d values were visualized in a heatmap. Refer to Supplemental Figure 5B for the standard error of the mean of each pair. **D**, Higher order relationships between vRNA segments were assessed with UPGMA dendrograms derived from mean d values. The point at which edges merge is equivalent to 1/2 *d*.

Genome-wide phylogenetic inference can distinguish parallel evolution of vRNA segments that interact during genomic assembly through RNA-RNA interactions in addition to that of the proteins encoded. To examine this, we compared all pairwise *d* values in H3N2 viruses from 2005-2014. **Figure 2C** shows the mean *d* values of all seven replicate trees for each pair of vRNA segments (refer to **Supplemental Figure 2B** for the standard error). To establish a threshold for significance in parallel evolution between vRNA trees, we determined a 95% confidence interval for *d* using a null dataset of randomly generated trees with an equivalent number of leaves (12 in this case) (**Supplemental Figure 3A & D**). Using this methodology, we found that low *d* values rarely occur by chance, with the vast majority of *d* values being greater than 15 in null trees. By comparison, mean *d* values ranged from 6.5 for PB1 & PA to 15 for PA & NS. Intriguingly, PB1, PA, NP, and NA were most highly coevolved with each other (all mean *d* values below 10), suggesting that parallel evolution between these four vRNA segments is enriched over other vRNA segments. Surprisingly, the PB2 tree was most congruent with the NA tree rather than the PB1 or PA trees, suggesting that the relationship between these segments may supercede the essential role of the PB2 protein in the polymerase complex. In contrast, the mean *d* values of the NS phylogenies with the four core vRNA segments were 14 to 15, approaching the 95% confidence threshold of 15.3. However, higher values of *d* may have resulted from a lack of statistical significance as neither the M or NS trees were strongly supported by bootstrapping (**Supplemental Figure 1B**). To further visualize higher order relationships between vRNA segments, we assembled a matrix of the pairwise *d* values between vRNA trees and constructed a dendrogram of the *d* values (**Figure 2D**). This dendrogram highlights our observation that parallel evolution is most robust between PB1, PA, NP and NA.

### Parallel evolution is largely conserved over time within H3N2 viruses

Recent studies have identified a highly plastic and redundant network of interactions between vRNA segments in influenza virions produced during productive infection, many of which may be transient (Dadonaite et al., 2019; Le Sage et al., 2020). Based on these observations, it is plausible that vRNA relationships defined by parallel evolution change over time. To examine the conservation of evolutionary relationships in H3N2 viruses, we estimated the mean and standard error of *d* for all pairs of vRNA trees within a set of H3N2 viruses from an earlier time period (1995-2004) (**Figure 3A** and **Supplemental Figure 2A**, respectively). As was seen in the H3N2 viruses from 2005-2014, the mean *d* values ranged widely from 4.5 to 14. PB1, PA, NP, and NA remained highly coevolved in this time period, with mean *d* values from 4.5 to 8. Statistical differences between *d* values from each time period were only found for the NS segment (*p*-adj < 0.05, **Figure 3C**). However, NS trees had consistently low bootstrap support (**Supplemental Figure 1**), so these differences may be attributable to insufficient resolution in the underlying trees. We constructed a dendrogram of the mean *d* values of the 1995-2004 trees to examine whether the hierarchical coevolution between vRNA segments observed in the 2005-2014 viruses was also apparent in the 1995-2004 viruses (**Figure 3B**). Comparison of these dendrograms (**Figures 2D** and **3B**) using the Robinson-Foulds distance revealed a *d* value of 6, which lies within the 95% confidence interval for a tree with eight leaves (**Supplemental Figure 3A-B**). Based on these data, we conclude that there was minimal difference in the phylogenetic relationships between vRNA segments in H3N2 viruses from these two time periods.

**Figure 3.**
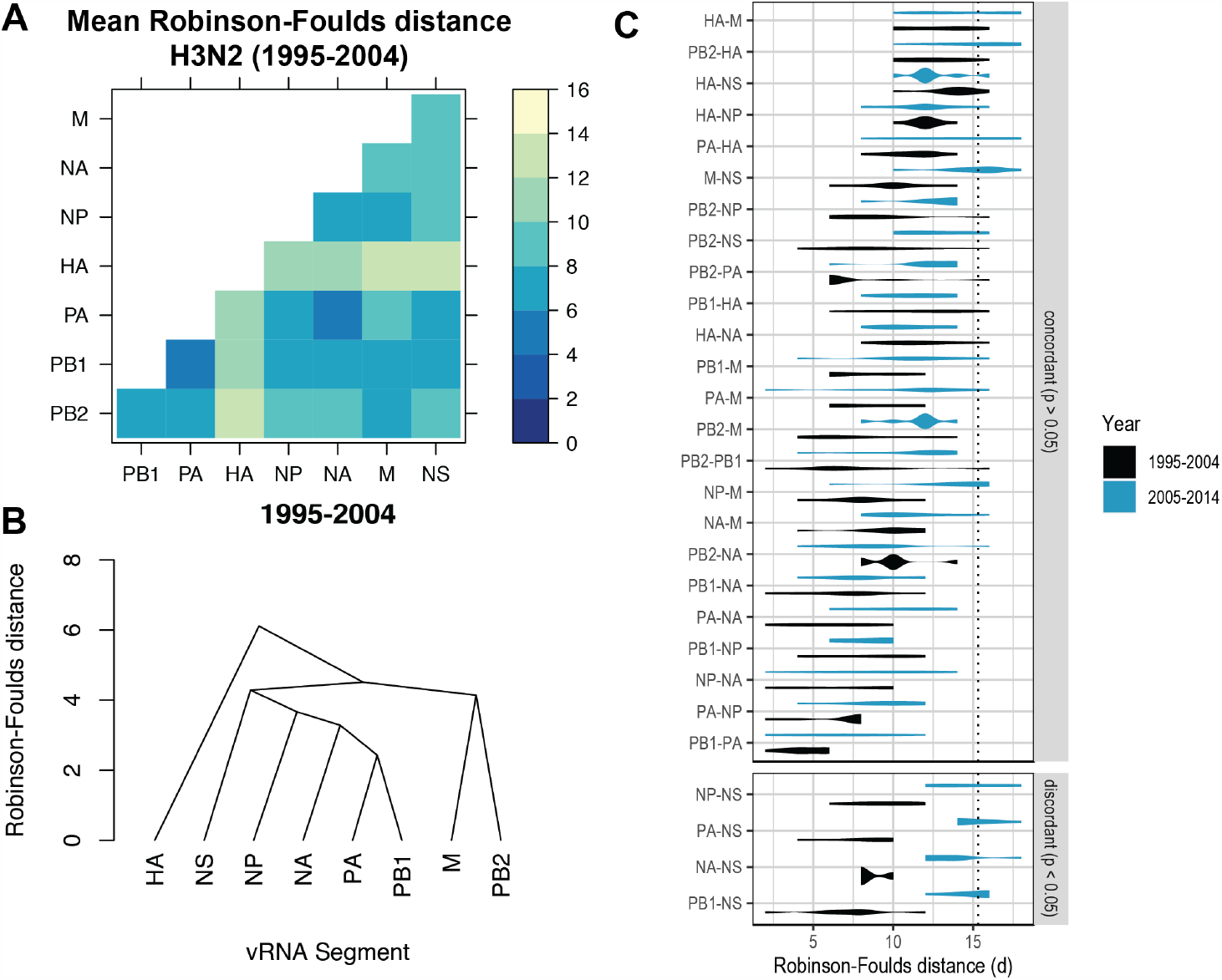
H3N2 virus vRNA coevolution is conserved through antigenic drift. **A**, Seven replicate maximum-likelihood phylogenies were reconstructed for each vRNA gene segment from human H3N2 virus sequences (1995-2004) as described in Figure 1. The Robinson-Foulds distance (*d*) between each pair of phylogenies was calculated for each replicate. The mean *d* values were visualized in a heatmap. Refer to Supplemental Figure 5A for the standard error of the mean of each pair. **B**, Higher order relationships between vRNA segments were assessed with UPGMA dendrograms derived from the mean *d* values. Scale bar corresponds to *d*. **C**, All seven replicate *d* values for each pair of trees were plotted comparing H3N2 viruses from 1995-2004 (black) to H3N2 viruses from 2005-2014 (turquoise). ‘Discordant’ pairs are clustered where *p <* 0.05 (Mann-Whitney *U* test with Benjamini-Hochberg correction). Dashed line, 95% confidence interval for phylogenetic concordance (determined by a null dataset; refer to Supplemental Figure 4).

### Evolutionary relationships between vRNA segments are dependent upon subtype and lineage

Our results suggest that vRNA relationships are remarkably consistent across H3N2 viruses from a period spanning two decades. To examine whether our approach captures anticipated changes in vRNA relationships, we assessed these relationships in H1N1 viruses from 2000-2008 and 2010-2018. H1N1 viruses from these time periods represent distinct lineages before and after the 2009 pandemic. This pandemic was caused by an antigenically shifted H1N1 virus that emerged from reassortment of two swine-origin viruses (Garten et al., 2009). Therefore, these two time periods represent distinct H1N1 virus lineages, and different evolutionary relationships between vRNA segments would be expected for each lineage.

Species trees comprising full-length concatenated H1N1 virus genomes from 2000-2008 or 2010-2018 were constructed and clusters were defined using the same approach described for H3N2 viruses. While this method produced a similar number of clusters for both sets of H1N1 viruses (**Table 1**), there were fewer clusters than in H3N2 viruses, owing to the higher rate of evolution observed in H3N2 viruses (Bedford et al., 2015). Seven replicate strains were selected from each cluster (**Supplemental Tables 3-4**) and vRNA trees were built (**Supplemental Figure 4**). **Figure 4A-B** shows the mean *d* values of all seven replicate trees for each pair of vRNA segments (refer to **Supplemental Figure 2C-D** for the standard error). The 95% confidence interval cutoff for Robinson-Foulds distances corresponding to trees with 9 leaves was 8.6 (**Supplemental Figure 3C**) and is the threshold used for statistical comparison of parallel evolution in vRNA segments from H1N1 strains.

**Figure 4.**
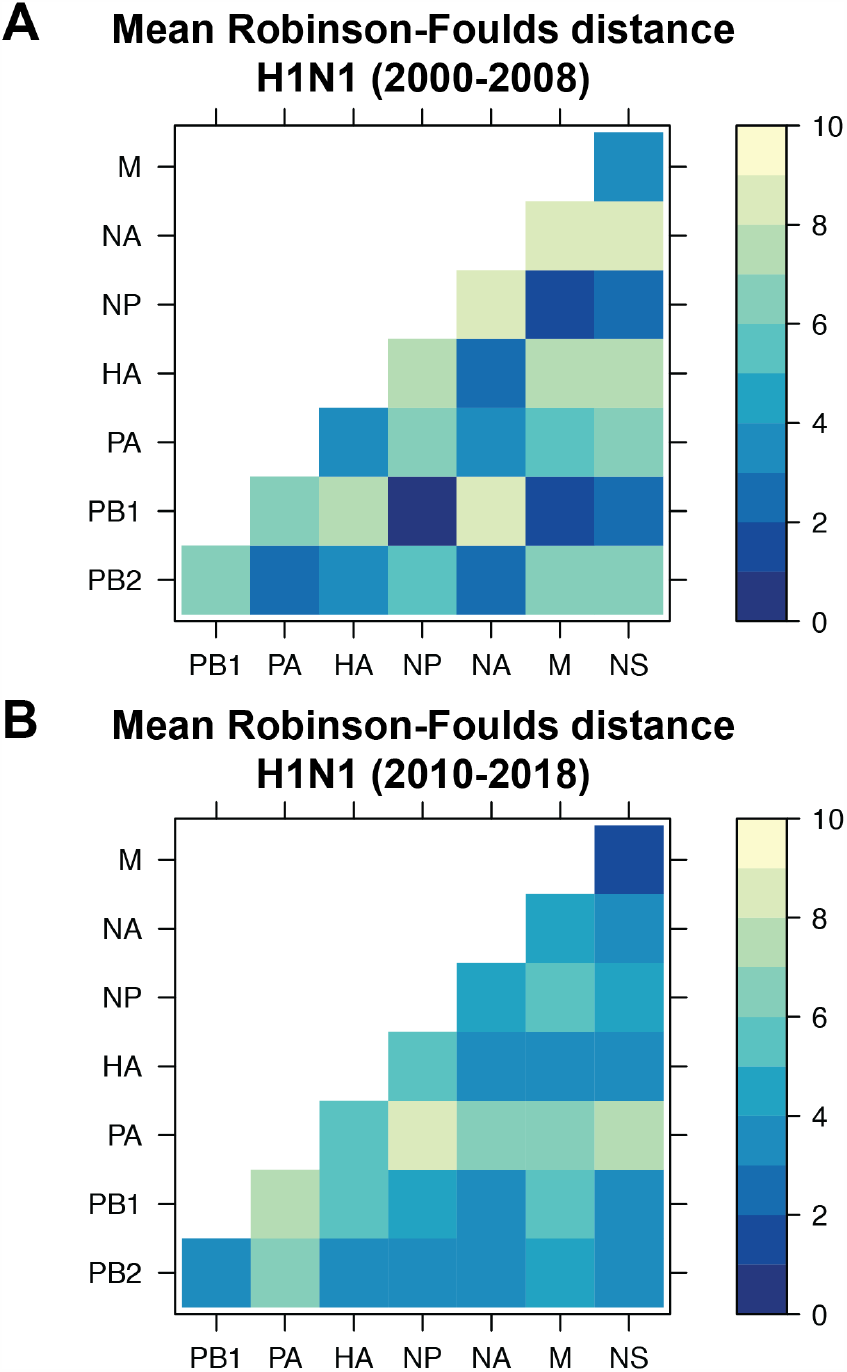
vRNA coevolution is dependent upon subtype and lineage. Seven replicate maximum-likelihood phylogenies were reconstructed for each vRNA gene segment from human H1N1 virus sequences from 2000-2008 **(A)** or 2010-2018 **(B)** as described in Figure 1. The pairwise Robinson-Foulds distance (*d*) between phylogenies in each replicate was calculated. The mean *d* values were visualized in a heatmap. Refer to Supplemental Figure 5C-D for the standard error of the mean of each pair.

Heatmaps comparing the *d* value between vRNA pairs suggest that vRNA relationships are not conserved across H1N1 viruses of different lineages (**Figure 4, A** vs **B**). We explored this further by constructing dendrograms of the *d* values for the pre-pandemic and post-pandemic H1N1 viruses (**Figure 5A-B**). The *d* value was 10 between pre-pandemic and post-pandemic H1N1 viruses, confirming that a high degree of incongruence existed between H1N1 viruses of different lineages (**Figure 5B**). To further explore individual differences between pairs of vRNA trees in H1N1 viruses of different lineages, we plotted *d* values for the pre-pandemic H1N1 viruses alongside the *d* values for the post-pandemic H1N1 viruses (**Figure 5C**). In stark contrast to the relatively conserved vRNA relationships observed in H3N2 viruses over time, many relationships between vRNA segments were disrupted in post-pandemic H1N1 viruses. Parallel evolution between PB1 and NP (mean *d* increased from 1 to 5; *p-adj* < 0.05, Mann-Whitney test) was notably displaced by stronger coevolution of PB1 with NA (mean *d* decreased from 9 to 3; *p-adj* < 0.05). The M and NS trees remained highly coevolved across H1N1 lineages, but each one was significantly more coevolved with the HA and NA trees in post-pandemic viruses (*p-adj* < 0.05). The PB2 trees diverged significantly from the PA trees in favor of greater parallel evolution with the NP and NS trees (*p-adj* < 0.05). Some of these data can be explained by weaker bootstrap support in H1N1 trees, particularly those from H1N1 viruses from 2010-2018 (**Supplemental Figure 4**). However, these data suggest that viruses from different lineages of the same subtype develop distinct vRNA evolutionary relationships, which is an important consideration when predicting reassortment between emerging viruses from different lineages.

**Figure 5.**
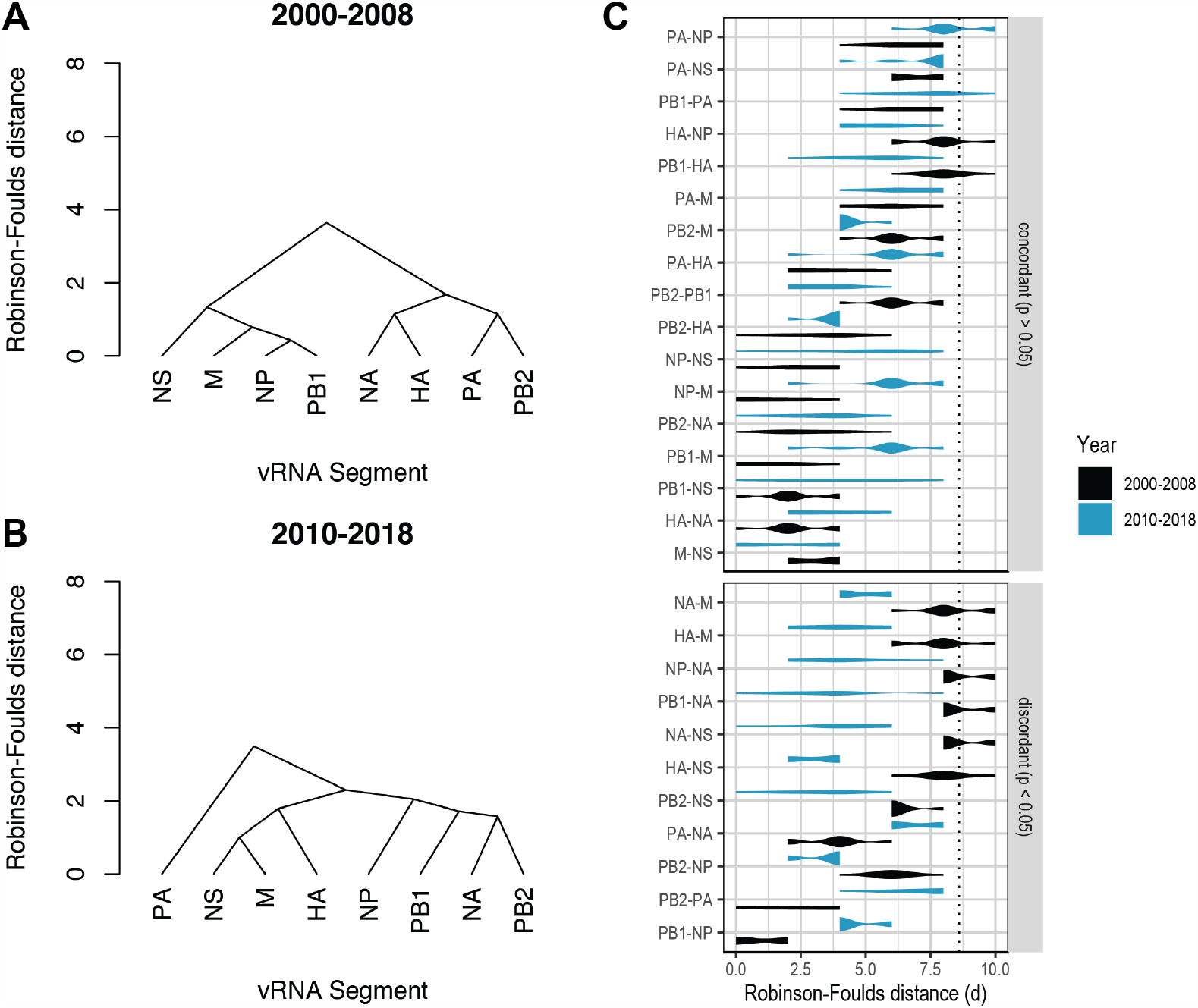
vRNA coevolution diverges in antigenically shifted H1N1 viruses. Higher order relationships between vRNA segments corresponding to H1N1 viruses from 2000-2008 **(A)** or 2010-2018 **(B)** were assessed with UPGMA dendrograms derived from the mean *d* values in Figure 4. Scale bar corresponds to *d*. **C**, All seven replicate *d* values for each pair of trees were plotted comparing H1N1 viruses from 2000-2008 (black) to H1N1 viruses from 2010-2018 (turquoise). ‘Discordant’ pairs are clustered where *p <* 0.05 (Mann-Whitney *U* test with Benjamini-Hochberg correction). Dashed line, 95% confidence interval for phylogenetic concordance (determined by a null dataset; refer to Supplemental Figure 4).

Comparison of the dendrograms from H1N1 and H3N2 viruses revealed some expected similarities as well as differences (**Fig 2D, 3D**, and **5A-B**). For example, PB1 and NP share a common evolutionary relationship across all four sets of influenza viruses examined in this study (**Figures 2D, 3B**, and **5A-B**), which may be expected based on the shared role of the encoded proteins in replication (Fodor, 2013). However, in H3N2 viruses PB1 and NP are next most closely related to NA and NP (**Figures 2D** and **3B**), while in H1N1 viruses their relationship with other segments varies (**Figure 5A-B**). We compared the overall similarity of all four dendrograms by computing the Robinson-Foulds distance. The pre-pandemic H1N1 virus dendrogram was significantly different from both of the H3N2 virus dendrograms: *d* = 10 when compared to either 1995-2004 or 2005-2014 H3N2 virus dendrograms. A *d* value of 10 is well outside of the 95% confidence interval cutoff of 6.5 determined by the null dataset for trees with 8 leaves (**Supplemental Figure 3A-B**). Likewise, *d* = 10 for the post-pandemic H1N1 virus dendrogram when compared to either of the H3N2 virus dendrograms. These results are in direct contrast to our previously determined distance for the H3N2 virus dendrograms to one another (*d* = 6). Overall, these data indicate that parallel evolution between vRNA segments is distinct between influenza subtypes isolated from humans within similar time scales.

### Parallel evolution in H3N2 viruses is not driven solely by protein-coding mutations

As discussed previously, a phylogenetics approach such as ours would encompass parallel evolution driven by either protein and RNA relationships. We have already shown that known protein relationships between PB1 and PA, two members of the polymerase complex, are identified by our approach (**Figure 2A**). However, the observation that PB2 is more coevolved with NA than with either PB1 or PA (**Figure 2C**) suggests that our method also reveals protein-independent parallel evolution, since these proteins are not known to function together during infection. Using H3N2 viruses from 2005-2014, which yielded vRNA trees with the highest overall bootstrap support (**Supplemental Figure 1B**), we explored the extent to which parallel evolution between vRNA segments is driven by protein-coding mutations. To do so, we converted the vRNA sequence alignments, which are negative-sense, into positive-sense RNA (i.e., coding sense) and translated the coding sequences into amino acid alignments. For the M and NS sequence alignments that encode two splice variants each, the M1/M2 and NS1/NS2 amino acid alignments were both translated and analyzed individually. Neighbor-joining trees were reconstructed from the amino acid alignments and the evolutionary relationships between H3N2 proteins were analyzed by the Robinson-Foulds distance. We took the resultant *d* values and constructed a dendrogram between all pairs of protein trees, as was previously done with vRNA trees (**Supplemental Figure 5**). This dendrogram appears distinct from the dendrogram built from the corresponding gene (vRNA) trees (**Figure 2D**). As might be expected, the greatest degree of parallel evolution lying at the core of this dendrogram was between HA and NA, two viral glycoproteins with coordinated functions in attachment, motility, and entry (Bloom et al., 2010; Sakai, Nishimura, Naito, & Saito, 2017).

To compare parallel evolution between influenza proteins to that of the parent vRNA segments, the mean *d* values from the gene trees were plotted against the mean *d* values from the protein trees (**Figure 6**). In the case of the M and NS segments, mean *d* values for all M1/M2 or NS1/NS2 combinations are shown. Many vRNA pairs, such as PB2 and PB1, lie along the identity line, indicating that protein interactions are more likely to drive parallel evolution in those vRNA segments. Interestingly, HA and NA were the only pair of vRNA segments that lay significantly above the identity line, strongly supporting the observation made by others that epistatic interactions between these proteins constrains their evolution (Jang & Bae, 2018; Kryazhimskiy et al., 2011; Neverov et al., 2015). Of particular interest was that several vRNA segments, such as PB2 and NA (**Figure 6, open diamond**), lay significantly below the identity line. This could be indicative of either purifying selection, or of greater parallel coevolution between the vRNA segments than the proteins encoded. While this is not altogether unexpected, considering that the mutation rate of a protein is unlikely to be as high as the mutation rate of the corresponding gene, we would expect conserved RNA interactions to also have this effect. These observations suggest that parallel evolution may identify putative RNA interactions between vRNA segments that could facilitate selective assembly and packaging.

**Figure 6.**
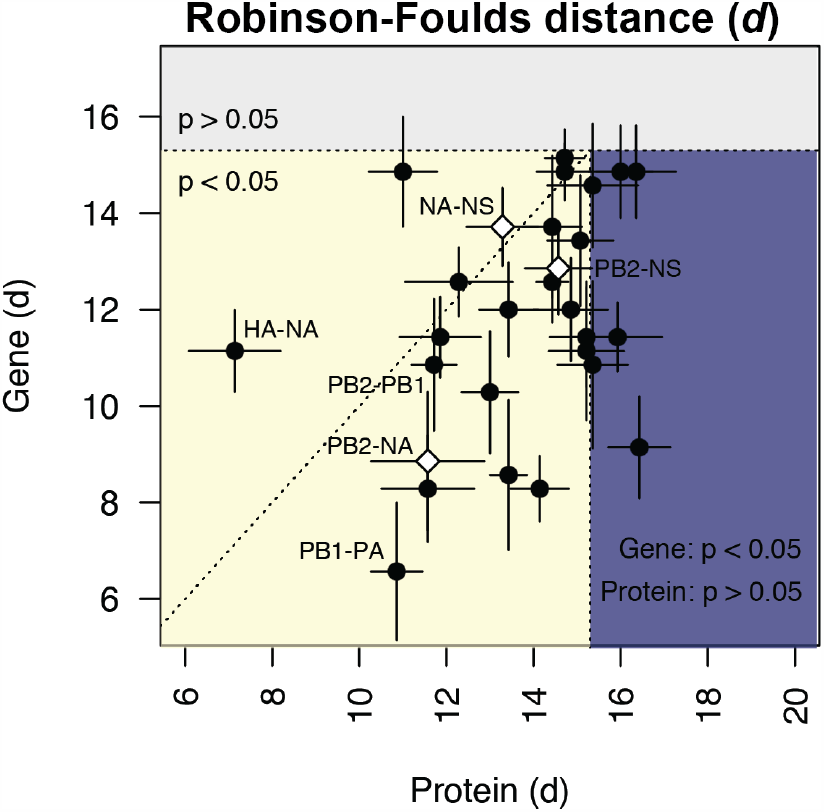
Protein-coding substitutions do not fully account for gene coevolution. H3N2 virus vRNA gene sequence alignments from 2005-2014 were translated into the corresponding amino acid alignments. Neighbor-joining phylogenies were reconstructed from these alignments and the Robinson-Foulds distance (*d*) was tabulated for all protein tree pairs. The mean *d* value of each pair of protein trees was plotted against the mean *d* value of the corresponding gene trees. For the M and NS gene segments, which encode multiple protein products, *d* values were calculated for each protein tree individually and the average *d* values were used. Error bars indicate the standard error of the mean (SEM) for replicate trees (n = 7). Dashed horizontal and vertical lines, 95% confidence interval (Cl) for phylogenetic concordance, as determined by a null dataset (refer to Supplemental Figure 4). The region shaded yellow lies within the 95% Cl for both gene and protein trees with the identity line plotted. The region shaded blue lies within the 95% Cl for gene trees but not protein trees. The region shaded gray lies outside the 95% Cl for both gene and protein trees.

### PB2 and NA viral ribonucleoprotein complexes (vRNPs) preferentially colocalize at the nuclear periphery in vitro

To address whether parallel evolution between the PB2 and NA segments corresponds with their behavior during influenza virus infection, we examined whether these vRNA segments preferentially colocalize in infected cells. During influenza virus infection, viral RNA are synthesized in the nucleus and then transported to the plasma membrane for packaging on endocytic vesicles (Lakdawala, Fodor, & Subbarao, 2016). We therefore examined intracellular colocalization of PB2, NA, and NS during infection with an H3N2 virus. These segments encompass a pair of segments with high gene-based parallel evolution (PB2-NA) as well as pairs with little evidence of parallel evolution (PB2-NS; NA-NS) (**Figures 2C** and **6, open diamonds**).

We quantified colocalization using our established method for examining intracellular colocalization of vRNA segments by fluorescence *in situ* hybridization (FISH) and immunofluorescence (IF) in productively infected cells (Lakdawala et al., 2014; Nturibi, Bhagwat, Coburn, Myerburg, & Lakdawala, 2017). Lung epithelial A549 cells were infected with an H3N2 virus representative of the time period analyzed (A/Perth/16/2009) for 8 hours and stained for three vRNA segments, NP protein, and nuclei. The NP antibody stain was used to normalize the pairwise colocalization data to the total number of vRNP foci present in the cells. Entire cell volumes were captured to analyze the colocalization of vRNA segments specifically within the cytoplasm by masking the signal within the nucleus. A representative image of an infected cell from one of three independently performed experiments is shown after deconvolution (**Figure 7A**) and segmentation of cytoplasmic foci (**Figure 7B**).

**Figure 7.**
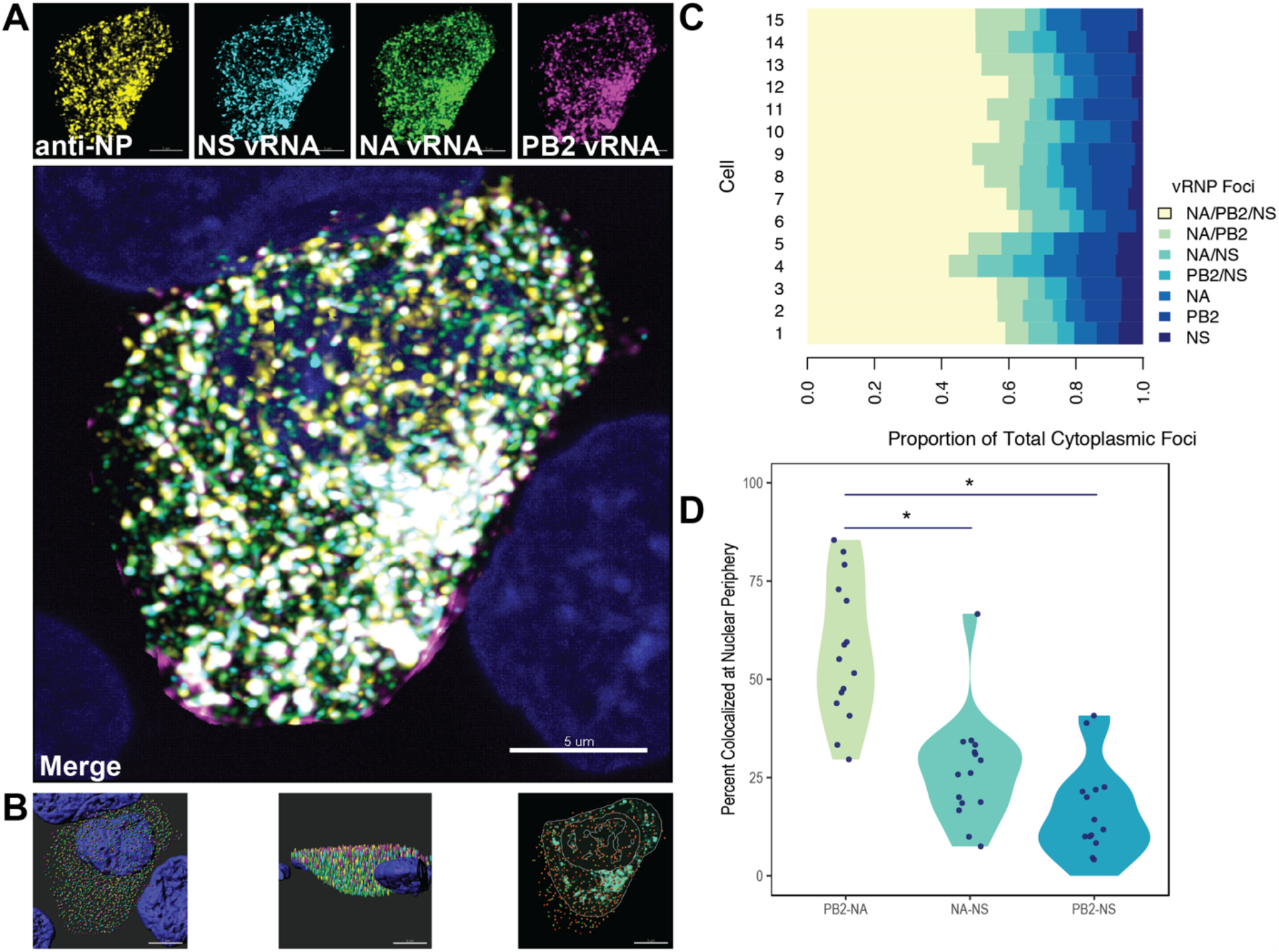
Colocalization of vRNA segments at the nuclear periphery correlates with evolutionary relationships during productive viral infection. A549 cells were infected with A/Perth/16/2009 (H3N2) at an MOI of 2 or mock infected. Cells were fixed at 8 hours post-infection and combination fluorescence *in situ* hybridization/immunofluoresence (FISH and IF, respectively) was performed. FISH probes targeting the NS, NA and PB2 vRNA segments were labeled with Alexa Fluor 488, Quasar 570 and Quasar 670, respectively. Antibodies targeting nucleoprotein (NP) were used with an anti-mouse Alexa Fluor 594 secondary antibody. Nuclei were labeled with DAPI. Coverslips were mounted and volumetric imaging was performed to obtain Nyquist sampling. **A**, A maximum projection image of a representative cell is shown after cell segmentation. Scale bar corresponds to 5 pm. **B**, A3D rendering of the cell after analysis. **C**, Colocalization of vRNA segments was assessed in 15 individual infected cells. **D**, Quantification of each pair of vRNA segments within 300 nm of the nuclear border. Each point represents an individual cell (n = 15). Aggregate data from three independently performed experiments are shown. * denotes *p <* 0.05 (Mann-Whitney *U* test)

Whole cytoplasmic analysis of vRNP foci colocalization in fifteen individually analyzed cells revealed that the majority of cytoplasmic foci contained all three NA, PB2 and NS segments (**Figure 7C**). This observation may represent the complex nature of genomic assembly, where all segments come together en route to the plasma membrane. In addition, the cytoplasm includes vRNA segments colocalized at the nuclear periphery prior to packaging, where there are more complete assembly complexes (Lakdawala et al., 2014). Given that newly synthesized vRNA segments are exported from the nucleus as assembly intermediates comprising greater than two but fewer than 8 segments, these perinuclear assembly intermediates may function as nodes for further assembly (Majarian, Murphy, & Lakdawala, 2018). Therefore, we assessed the potential for PB2, NA and NS to colocalize at the nuclear periphery, where assembly intermediates first begin to form. We defined localization at the nuclear periphery to within three-hundred nanometers, the limit of resolution in this system. Examination of newly exported vRNP complexes within three-hundred nanometers of the nuclear periphery revealed an enrichment of PB2-NA vRNP complexes over either NA-NS or PB2-NS vRNP complexes. These data suggest that PB2 and NA vRNA segments could comprise one such node from which to build the entire complex of all eight segments and support the hypothesis that RNA assembly interactions, in addition to protein interactions, can drive parallel evolution between vRNA segments in influenza viruses.

## Discussion

In this study, we used phylogenetics and molecular biology methods to investigate genome-wide relationships between vRNA segments in seasonal human influenza A viruses. We found that parallel evolution varies considerably between vRNA segments, with distinct relationships forming in different influenza virus subtypes (H1N1 vs H3N2) and between H1N1 virus lineages that arose from distinct host origins. We further demonstrate that evolutionary relatedness between vRNA segments in H3N2 viruses is largely conserved over time. Importantly, our data suggest that parallel evolution cannot be attributed solely to protein interactions, and we successfully predicted intracellular colocalization between two coevolved vRNA segments during infection with an H3N2 virus. Thus, we present a phylogenetic approach for interrogating putative RNA associations that could be broadly applied toward the study of genomic assembly and reassortment in segmented viruses.

Selective assembly of all eight genomic segments is fundamental to the production of fully infectious virus particles. We and others have used a variety of biochemical approaches to investigate the mechanisms that promote selective assembly (Dadonaite et al., 2019; Le Sage et al., 2020). We previously demonstrated that binding of vRNA segments by the NP protein is non-uniform and non-random (Le Sage et al., 2018; Lee et al., 2017), supporting the model that intersegmental RNA interactions facilitate selective assembly. Biochemical approaches to define bona fide intersegmental RNA-RNA interactions demonstrated that the interaction network is highly flexible and varies between H1N1 and H3N2 viruses (Dadonaite et al., 2019; Le Sage et al., 2020). These observations are consistent with our conclusion that RNA interactions constrain parallel evolution between vRNA segments in a manner sensitive to the genetic context studied.

The approach we present here differs from other experimental approaches in that we identify a novel, conserved RNA-based relationship in H3N2 viruses. For example, we found that relationships between PB1, PA, NP and NA are enriched over other segments in H3N2 viruses and conserved over time. One might expect PB1, PA and NP to coevolve because of the functions of the proteins they encode: the polymerase subunits PB2, PB1 and PA form a supramolecular complex around each vRNA segment with NP (Fodor, 2013). However, this explanation does not account for the parallel evolution observed between vRNP protein components and NA, and our microscopy data demonstrates that the NA segment preferentially colocalizes with the vRNA of one such vRNP component, supporting the possibility that parallel evolution of NA with PB1, PA and NP could also be driven by RNA interactions. These observations suggest that RNA relationships with the NA segment may facilitate selective assembly of vRNA segments. Further work should be directed at determining the underlying nature driving the novel relationship between these segments.

Previous pandemic influenza viruses emerged through reassortment (Neumann et al., 2009). Prediction of future influenza pandemics relies on understanding assembly of vRNA segments within a cell. As we have discussed, experimental investigations of intersegmental RNA interactions indicate that the vRNA interactome is distinct among virus strains and highly plastic (Dadonaite et al., 2019; Le Sage et al., 2020). Therefore, experimental approaches are unlikely to provide the holistic view necessary to predict reassortment outcomes of two circulating influenza strains. In contrast, we identified several conserved relationships between vRNA segments in H3N2 viruses that could impose constraints on reassortment. Thus, investigation of epistatic relationships between vRNA segments through phylogenetics could inform the sequence-based prediction of barriers to reassortment in emerging influenza viruses.

## Materials and Methods

### Viruses and cells

Human adenocarcinoma alveolar basal epithelial cells (A549, ATCC) were maintained in high-glucose Dulbecco’s Modified Eagle Medium (DMEM, Sigma) supplemented with 10% fetal bovine serum (FBS, HyClone), 2% L-glutamine, and 1% penicillin/streptomycin. Madin-Darby canine kidney epithelial cells (MDCK, ATCC) were maintained in MEM supplemented with 10% FBS. All cells were maintained at 37°C in 5% CO^2^. Reverse genetics plasmids of the influenza A virus, A/Perth/16/2009 (H3N2), were kindly provided by Dr. Jesse Bloom (Fred Hutchinson Cancer Research Center, Seattle). Recombinant virus was rescued as previously described (Lakdawala et al., 2011). Virus titers were determined by 50% tissue culture infectious dose (TCID^50^) on MDCK-SIAT cells using the endpoint titration method (Reed & Muench, 1938).

### Influenza A virus sequences

FASTA files of each genomic segment of human influenza A virus sequences of H1N1 and H3N2 viruses were downloaded from the Influenza Research Database (IRD, http://www.fludb.org) (Zhang et al., 2016) on June 22, 2018 and July 3, 2018, respectively. Strains lacking full-length genomic sequence data were excluded.

### Clustering and sequence selection

Sequences were read into R (version 3.5.2) using the DECIPHER (version 2.18.1) package (Wright, 2015) and subset into the time periods 1995-2004 and 2005-2014 (H3N2 strains) or 2000-2008 and 2010-2018 (H1N1 strains). Time periods were selected in part to ensure a similar level of genetic diversity between strains. In each strain, all eight vRNA segments were concatenated into a full-length genome from which alignments were constructed. A neighbor-joining species tree was built by clustering strains into operational taxonomic units with sequence identity cutoffs ranging from 95-99%. In H3N2 viruses from 1995-2004, there were 3, 7, 16, 53, and 259 clusters corresponding to cutoffs of 95%, 96%, 97%, 98%, and 99% sequence identity, respectively. The 95-96% sequence identity cutoffs were discarded, as these produced trees with an insufficient number of branches for comparison by the Robinson-Foulds distance. However, as the cutoff for sequence identity was increased from 97% to 99%, we observed a corresponding decrease in bootstrap support for trees built from representative sequences. A sequence identity cutoff of 97% was therefore selected to ensure the greatest degree of robustness in tree topologies. Small clusters occurred infrequently and were either omitted or collapsed into a single cluster. Seven replicate strains were randomly chosen from each cluster for further study and visually inspected for sequencing ambiguities. A list of all strains analyzed and the corresponding accession numbers can be found in **Supplemental Tables 1-4**.

### Phylogenetic tree reconstruction and analysis

Maximum-likelihood phylogenies were reconstructed under an HKY85 model for either full-length genomes or individual vRNA segments with 100 or 1,000 bootstrap replicates, as indicated, using the DECIPHER package in R. The phangorn package (version 2.5.5) (Schliep, 2011) was used to identify an appropriate model of evolution for phylogenetic reconstruction. Strain names are coded by cluster number in all trees. Phylogenetic trees of full-length concatenated genomes are shown in **Supplemental Figure 6**. Neighbor-joining protein phylogenies were built from amino acid alignments after translation of the corresponding coding sequence alignments.

Tanglegrams were built from pairs of vRNA phylogenies within replicates using the phytools package (version 0.7-70) (Revell, 2012). The Robinson-Foulds distance (*d*) was calculated for each pair of phylogenies using the ape package (version 5.4-1) (Paradis & Schliep, 2019).

Dendrograms visualizing the mean *d* values between vRNA segments were built using the UPGMA method. Tanglegrams were constructed between each pair of dendrograms and a *d* value was calculated for each tanglegram.

### Fluorescence in situ hybridization and immunofluorescence

Custom Stellaris RNA FISH oligonucleotide probes specific for the H3N2 virus NS, NA and PB2 vRNA segments were purchased from BioSearch Technologies (refer to **Supplemental Table 5** for FISH probe sequences). Each custom probe mix is comprised of 20 to 40 20-mers that span the length of the vRNA segment of interest. Probes with high complementarity against other vRNA segments or positive-sense RNA were excluded during the design process. The NS probe was purchased with a terminal amine group and manually conjugated to the Alexa Fluor 488 fluorophore using the Alexa Fluor 488 Oligonucleotide Amine Labeling Kit (Invitrogen). The NA and PB2 probes were labeled by the manufacturer with the Quasar 570 and Quasar 670 fluorophores, respectively.

Three independent fluorescence *in situ* hybridization and immunofluorescence (FISH-IF) experiments were performed. A549 cells were seeded directly onto 1.5 mm circular coverslips (Fisher Scientific) in tissue culture dishes. The next day, cells were infected at a multiplicity of infection (MOI) of 2 with A/Perth/16/2009 (H3N2) or mock infected in diluent. Cells were fixed at 8 hours post-infection with 4% paraformaldehyde and permeabilized overnight in ice cold 70% ethanol. Prior to hybridization, cells were rehydrated in wash buffer (10% formamide and 2X saline sodium citrate [SSC] in DEPC-treated H^2^O) and then incubated at 28°C overnight in hybridization buffer (10% dextran sulfate, 2 mM vanadyl-ribonucleoside complex, 0.02% RNA-free BSA, 1 mg/ml *E. coli* tRNA, 2X SSC, and 10% formamide in DEPC-treated H^2^O) with anti-influenza A virus NP antibody (Millipore, 1:2,000) and FISH probes. After hybridization, cells were washed and incubated with Alexa Fluor 594 goat anti-mouse (Invitrogen, 1:2,000) and DAPI (Sigma, 1:5,000) in wash buffer. Coverslips were mounted on slides in ProLong Diamond antifade mountant (Thermo Fisher).

### Confocal Imaging

Microscope slides were imaged on a Leica SP8 confocal microscope equipped with a pulsed white light laser as an excitation source and an acousto-optical beam splitter (AOBS) and Leica Hybrid Detectors. All imaging was performed with a 100X oil immersion objective with a numerical aperture of 1.4. Sequential scanning with a line averaging of 3 between frames was used. To obtain Nyquist sampling, z-stacks of each cell were taken with a step size of 0.17 µm to achieve a pixel size of 45 nm × 45 nm × 170 nm. The following custom parameters were established using single-color infected controls for sensitive detection of all five fluorophores: 405 nm excitation wavelength (λ_ex_) with 0.5% laser power and a detection range of 415 to 470 nm (PMT1; DAPI), 488 nm λ_ex_ with 10% laser power and a detection range of 493 to 540 nm with time gating of 1 to 6 nanoseconds (ns) (HyD4; Alexa Fluor 488), 582 λ_ex_ with 15% laser power and a detection range of 590 to 635 nm with time gating of 1.5 to 6 ns (HyD4; Cal Fluor Red 590), 545 nm λ_ex_ with 5% laser power and a detection range of 545 to 568 nm with time gating of 1.5 to 6 ns (HyD4; Quasar 570), 647 nm λ_ex_ with 5% laser power and a detection range of 670 to 730 nm with time gating of 1.5 to 6 ns (HyD5; Quasar 670). In each experiment, five volumetric z-stacks were imaged of infected cells and one z-stack was imaged of mock infected cells.

### FISH Colocalization

Background subtraction and deconvolution of confocal images were performed manually for each channel using Huygens Essential software (version 19.04, Scientific Volume Imaging B.V.). In each experiment, images taken of mock-infected cells were deconvolved using the same parameters as those of infected cells. 3D reconstruction and colocalization analysis of the resulting images were performed using Imaris software (version 8.4.2, Bitplane AG) as previously described (Lakdawala et al., 2014; Nturibi et al., 2017). Briefly, the cell of interest in each image was segmented using the ‘Surfaces’ and ‘Cell’ tools in Imaris software. DAPI signal was used to mask nuclear signal from the remaining channels. The ‘Spots’ tool was then used to populate the reconstructed cell with four different sets of Spots corresponding to foci from each of the remaining channels. In each experiment, the mock infected cell was analyzed in an identical manner and the fluorescence intensity for each channel of the mock-infected cell was used to establish fluorescence intensity thresholds at which 97% or more of the background signal was removed prior to Spot generation. A modified Matlab extension was then used to quantify spot colocalization using a distance threshold of 300 nm as previously described (Nturibi et al., 2017). Colocalization data was imported into the Cell and all data was exported and analyzed in R.

### Statistics

Sets of null trees were used to determine confidence intervals for the Robinson-Foulds distance (*d*) between phylogenetic trees. A set of 1,000 randomly sampled, unrooted trees with 8, 9, or 12 tips were built using the ape package. The Robinson-Foulds distance (*d*) was calculated for all pairs of trees and these were fit to a linear regression model. Null *d* values were either log-transformed or transformed by the Yeo-Johnson method (Yeo & Johnson, 2000), as indicated. Mean *d* values calculated for pairs of vRNA trees were considered significant if they fell within the first five percentiles as compared to null *d* values from random trees with the same number of tips (i.e. 95% of null *d* values were greater than the mean *d* value for a given pair of vRNA trees). A Mann-Whitney *U* test with a Benjamini-Hochberg post-hoc correction was used to identify statistically significant differences between *d* values from two time periods. A Mann-Whitney *U* test was also used to determine statistical significance of FISH-IF colocalization data.

## Code availability

Custom code for analysis of parallel evolution in concatenated, full-length genomic influenza virus sequences is available on GitHub (https://github.com/Lakdawala-Lab/Parallel-Evolution/).

## Supplemental Material

**Supplemental Figure 1.**
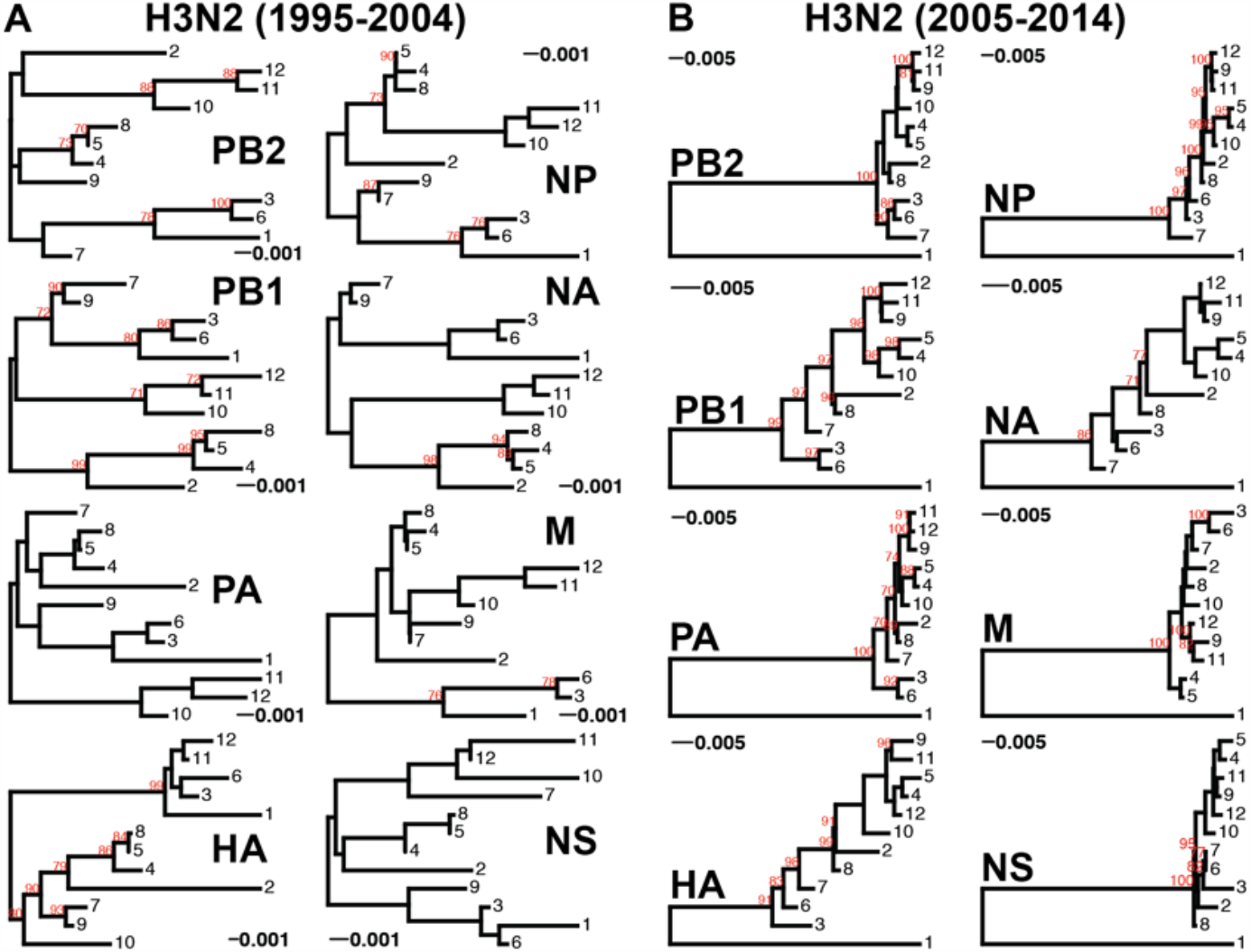
Phylogenies of vRNA segments from H3N2 virus sequences. Maximum-likelihood phylogenetic reconstruction of each vRNA segment from H3N2 virus sequences shown in Supplemental Tables 1-2. **A**, Representative phylogenies from replicate 2 for the 1995-2004 viruses. **B**, Representative phylogenies from replicate 7 for the 2005-2014 viruses. Sequences are coded by OTU cluster. Bootstrapping was performed with 100 replicates (bootstrap values greater than 70 are shown in red). Scale bars indicate percent divergence.

**Supplemental Figure 2.**
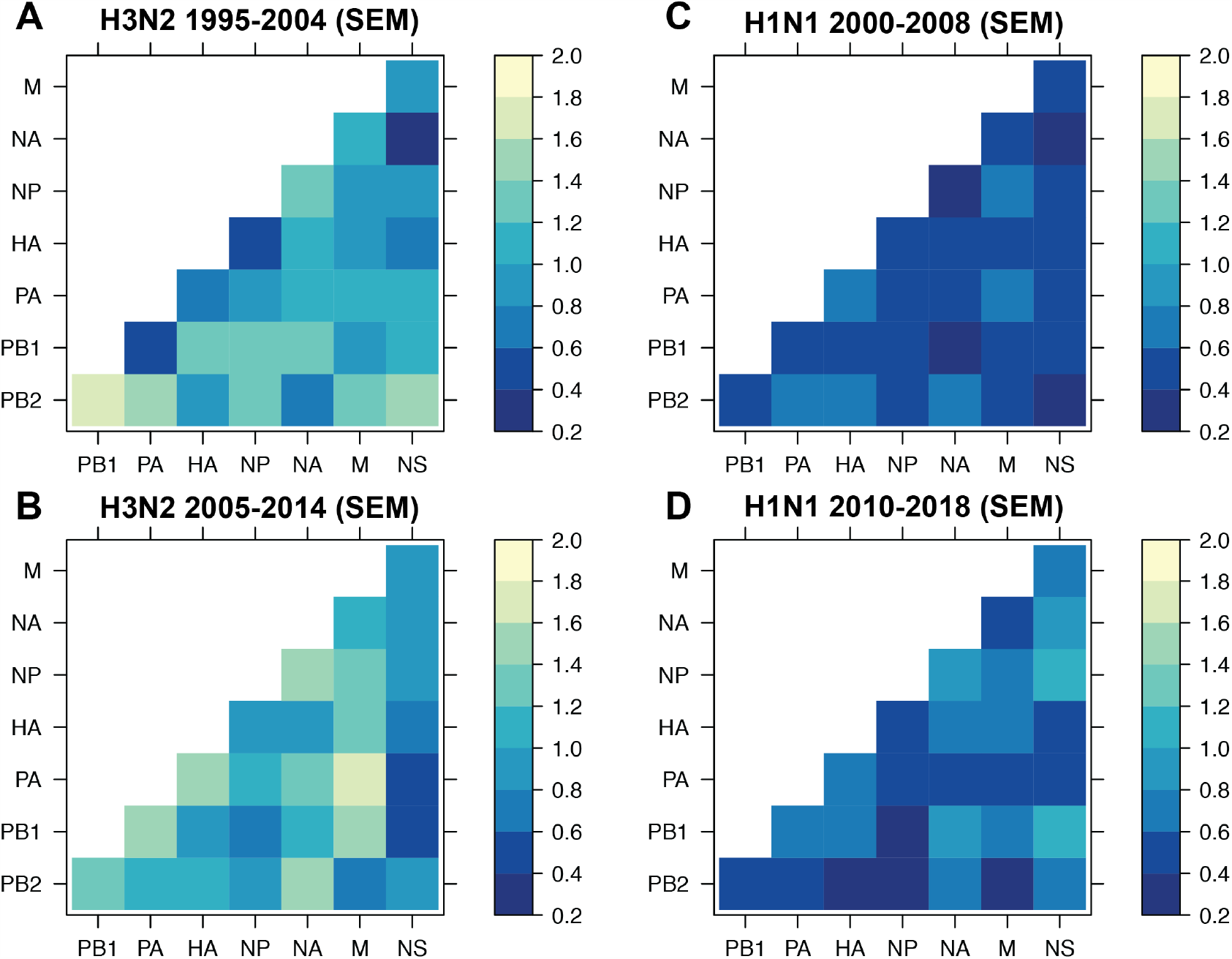
The standard error of the mean (SEM) of replicate Robinson-Foulds distances. The SEM of all pairwise Robinson-Foulds distances (*d*) was determined for H3N2 viruses from 1995-2004 (corresponding mean *d* values shown in Figure 3A) **(A)**, H3N2 viruses from 2005-2014 (corresponding mean *d* values shown in Figure 2C) **(B)**, H1N1 viruses from 2000-2008 (corresponding to mean *d* values from Figure 4A) **(C)**, and H1N1 viruses from 2010-2018 (corresponding to Figure 4B) **(D)**.

**Supplemental Figure 3.**
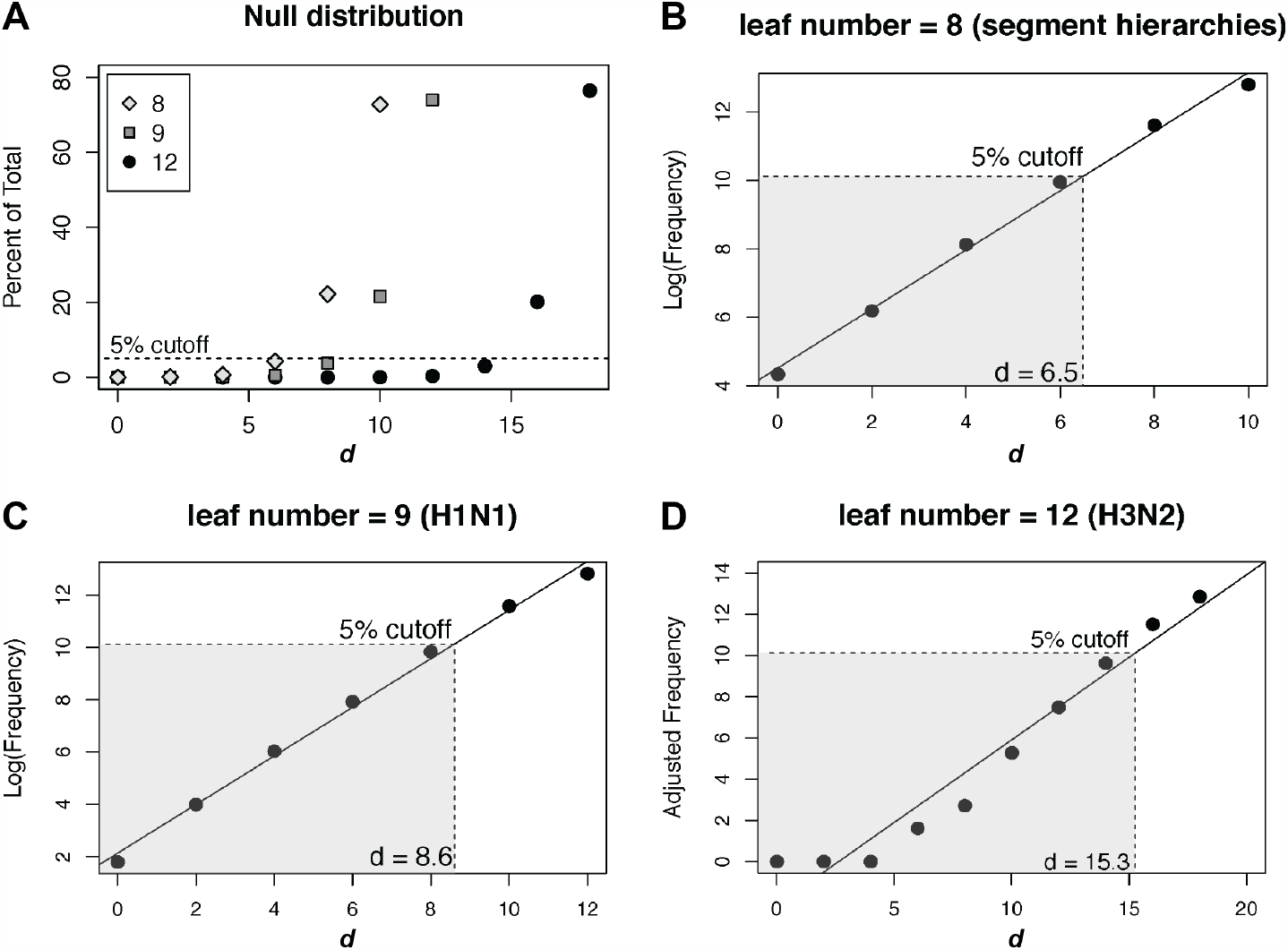
Null distribution of d values. **A**, The null distribution of *d* values in sets of 1,000 randomly sampled, unrooted trees with either 8 (gray diamonds, corresponding to the number of tips in each hierarchical dendrogram), 9 (gray squares, the number of tips in each H1N1 phylogeny), or 12 (black circles, the number of tips in each H3N2 phylogeny) leaf tips was determined. A dashed line demarcates the threshold for the 95% confidence interval. **B-D**, The null distributions shown in **A** were log transformed **(B, C)** or Yeo-Johnson transformed (Yeo & Johnson, 2000) **(D)** and fit to a linear regression model to establish the cutoff for the first five percentiles (shaded region), which was set as the 95% confidence interval cutoff.. The value of *d* at which 95% confidence is exceeded is indicated.

**Supplemental Figure 4.**
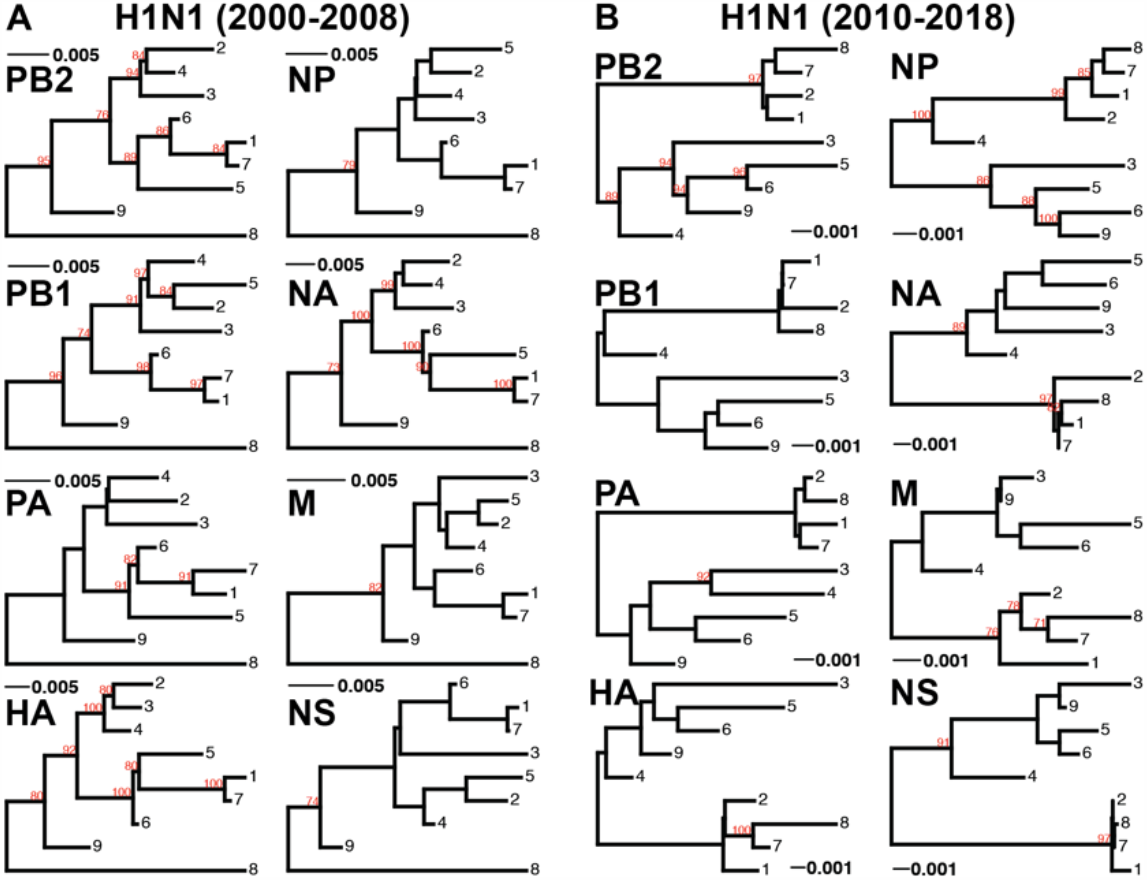
Phylogenies of vRNA segments from H1N1 virus sequences. Maximum-likelihood phylogenetic reconstruction of each vRNA segment from H1N1 virus sequences shown in Supplemental Tables 3-4. **A**, Represent ative phylogenies from replicate 6 for the 2000 -2008 viruses. **B**, Representative phylogenies from rep licate 3 for the 2010-2018 viruses. Sequences are coded by OTU cluster. Bootstrapping was performed with 100 replicates (bootstrap values greater than 70 are shown in red). Scale bars indicate percent divergence.

**Supplemental Figure 5.**
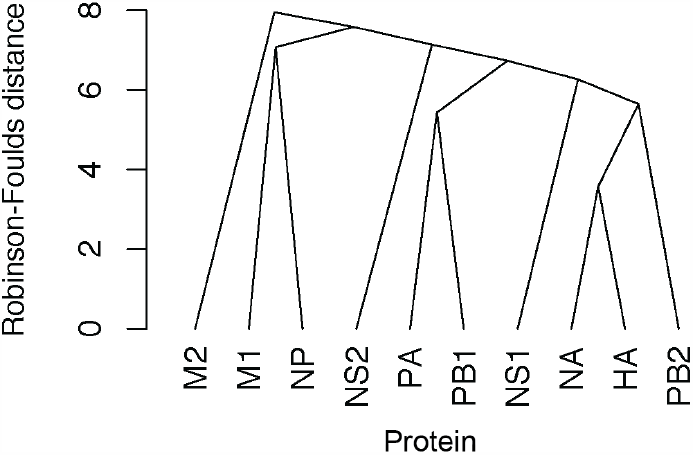
Coevolution of influenza A virus proteins in H3N2 viruses from 2005-2014. H3N2 virus vRNA gene sequence alignments were translated into the corresponding amino acid alignments. Neighbor-joining trees were reconstructed from these alignments and the Robinson-Foulds distance (*d*) was tabulated for all protein tree pairs. Higher order relationships between IAV proteins were assessed in a UPGMA dendrogram. Scale bar corresponds to *d*.

**Supplemental Figure 6.**
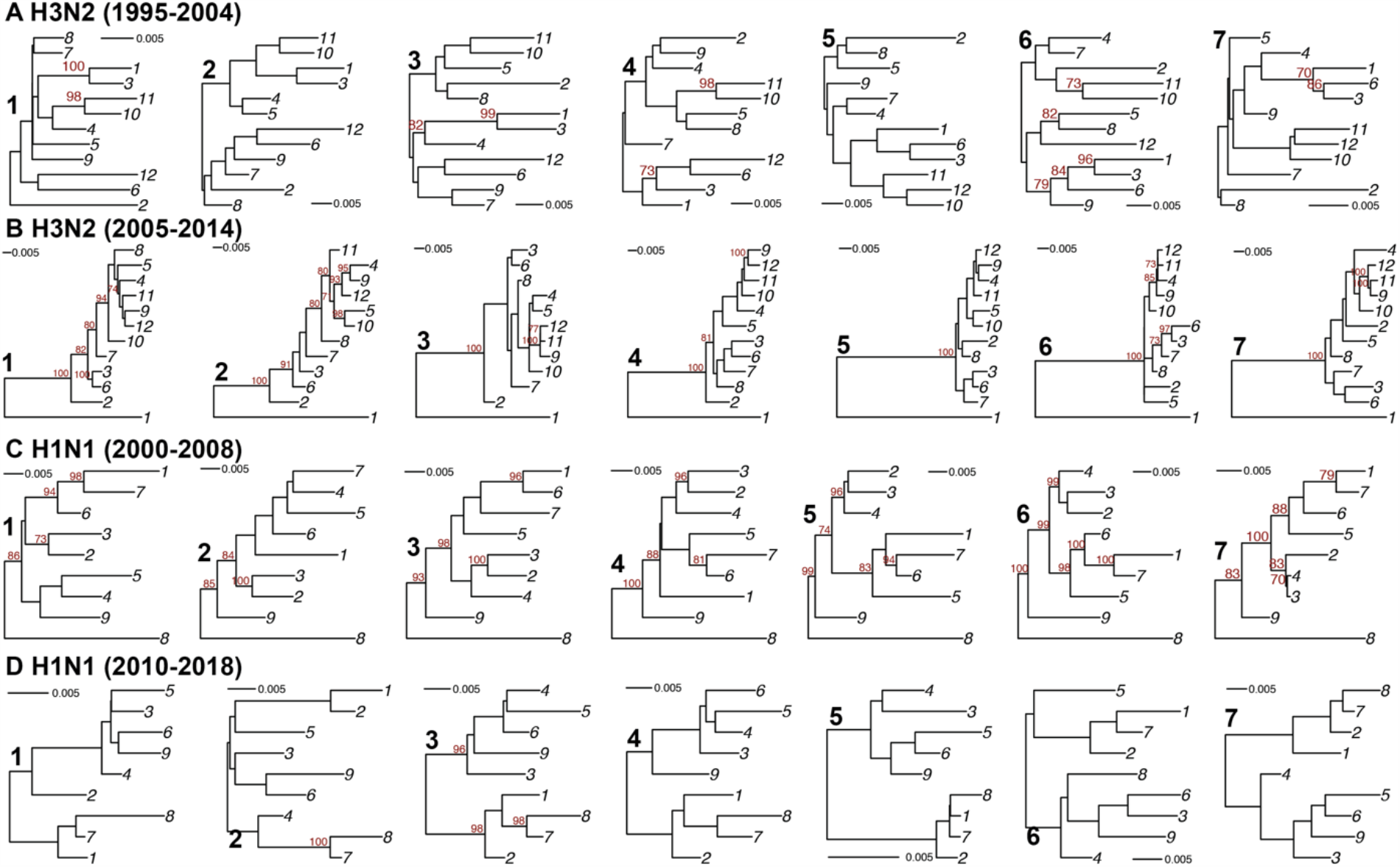
Replicate phylogenies of full-length concatenated H3N2 and H1N1 virus sequences. Maximum-likelihood phylogenetic reconstruction of full-length H3N2 virus (**A-B**) and H1N1 virus (**C-D**) genomic sequences shown in Supplemental Tables 1-4. Sequences are coded by OTU cluster. The numbers 1-7 in bold indicate replicates. Bootstrapping was performed with 1,000 replicates (bootstrap values greater than 70 are shown in red). Scale bars indicate percent divergence.

**Supplemental Table S1.**
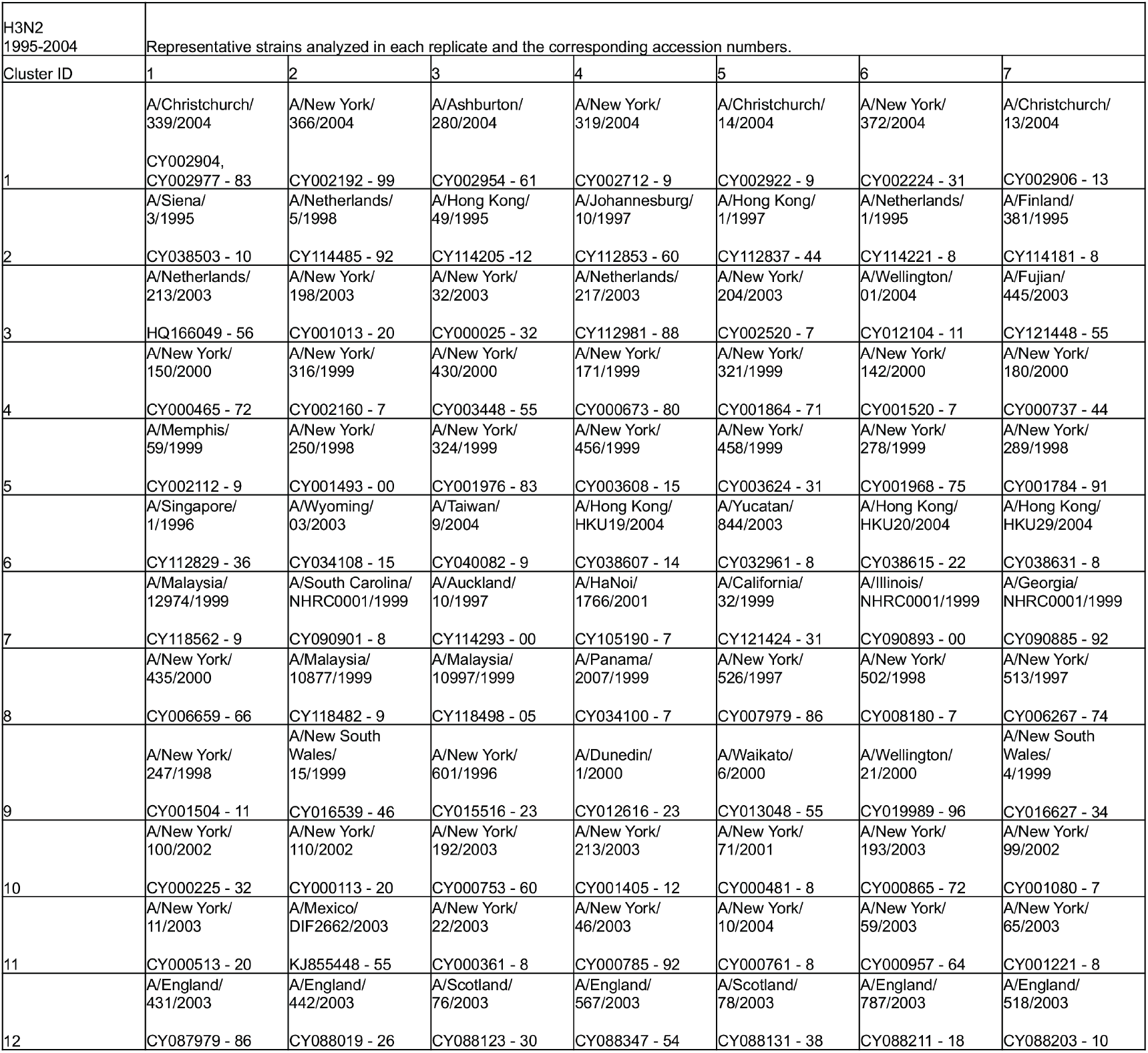
Human H3N2 sequences analyzed from 1995-2004 and the corresponding GenBank accession numbers. Human H3N2 sequences from 1995-2004 were downloaded from the Influenza Research Database and full-length genomes were concatenated and grouped into operational taxonomic units (numbered 1-12 under Cluster ID) with at least 97% sequence identity. Representative sequences were selected from these clusters for further analysis. Each vertical column indicates one replicate (seven replicates total).

**Supplemental Table S2.**
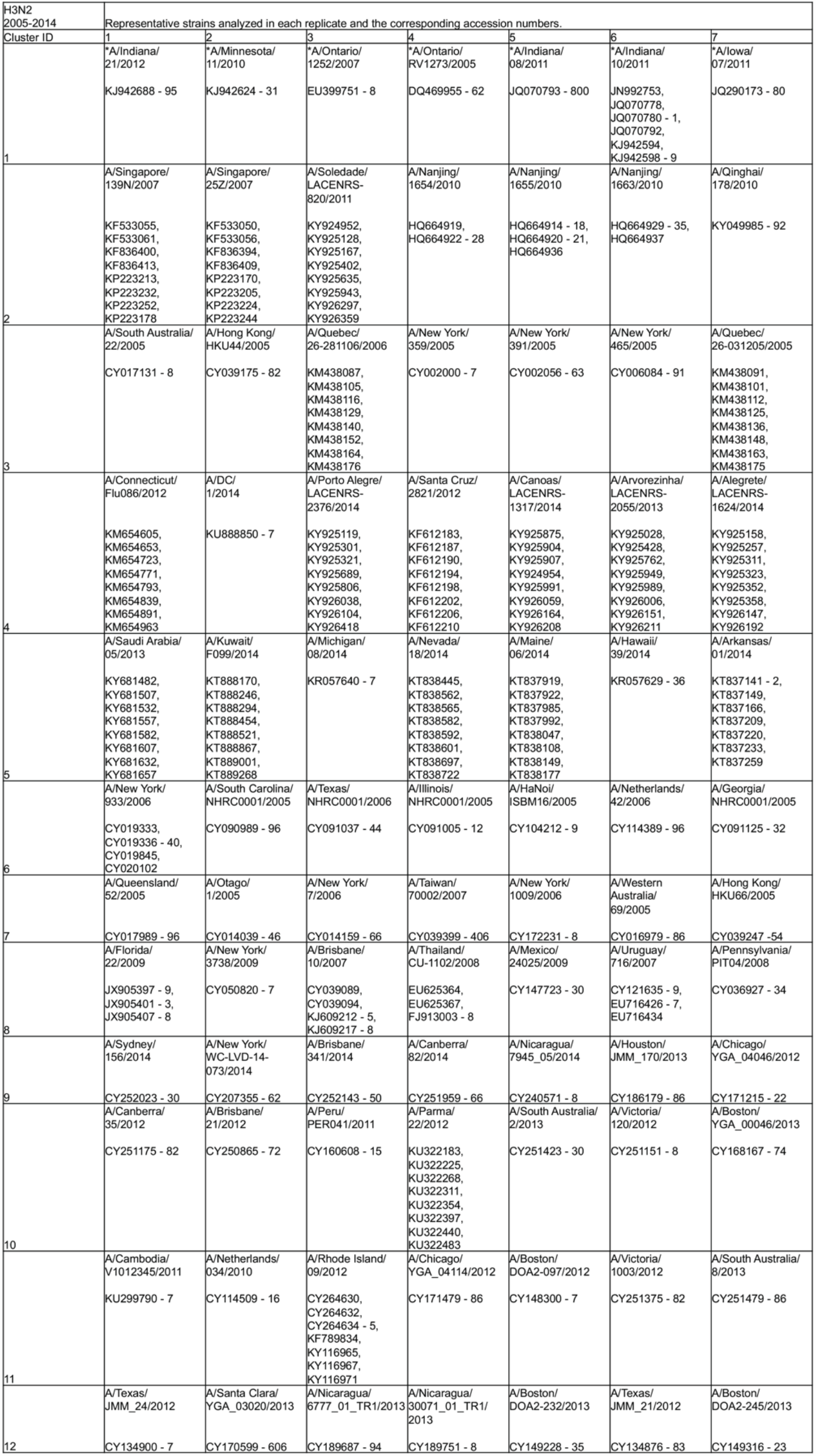
Human H3N2 sequences anal yzed from 2005-2014 and the corresponding GenBank accession numbers. Human H3N2 sequences from 2005-2014 were downloaded from the Influenza Research Database and full-length genomes were concatenated and grouped into operational taxonomic units (numbered 1-12 and labeled ClusterID) with at least 97% sequence identity. Representative sequences were selected from these clusters for further analysis. Each vertical column indicates one replicate (seven replicates total). Asterisks denote H3N2v strains.

**Supplemental Table S3.**
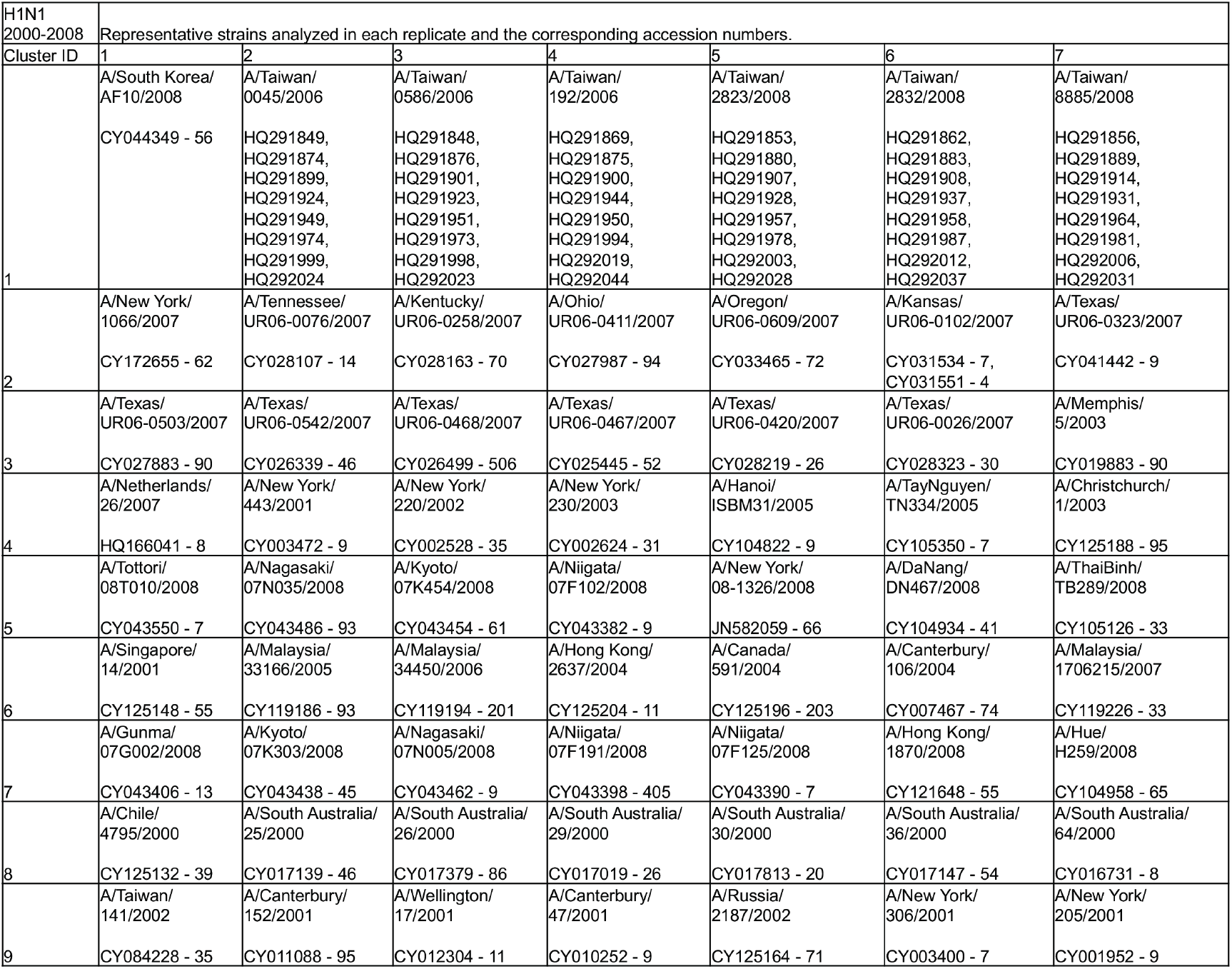
Human H1N1 sequences analyzed from 2000-2008 and the corresponding GenBank accession numbers. Human H1N1 sequences from 2000-2008 were downloaded from the Influenza Research Database and full-length genomes were concatenated and grouped into operational taxonomic units (numbered 1-9 under Cluster ID) with at least 97% sequence identity. Representative sequences were selected from these clusters for further analysis. Each vertical column indicates one replicate (seven replicates total).

**Supplemental Table S4.** Human H1N1 strains analyzed from 2010-2018 and the corresponding GenBank accession numbers. Human H1N1 sequences from 2010-2018 were downloaded from the Influenza Research Database and full-length genomes were concatenated and grouped into operational taxonomic units (numbered 1-9 under Cluster ID) with at least 97% sequence identity. Representative sequences were selected from these clusters for further analysis. Each vertical column indicates one replicate (seven replicates total).

| H1N1<br>2010-2018 | Representative strains analyzed in each replicate and the corresponding accession numbers. |  |  |  |  |  |  |
| --- | --- | --- | --- | --- | --- | --- | --- |
| Cluster ID | 1 | 2 | 3 | 4 | 5 | 6 | 7 |
| 1 | A/Gramado/<br>LACENRS-1287/2016<br>KY925104,<br>KY925179,<br>KY925188,<br>KY925751,<br>KY925798,<br>KY925960,<br>KY926084,<br>KY926299 | A/Taipei/<br>0045/2016<br>KY487178,<br>KY487296,<br>KY487367,<br>KY487426,<br>KY487456,<br>KY487565,<br>KY487602,<br>KY487714 | A/Vacaria/<br>LACENRS-495/2016<br>KY925355,<br>KY925365,<br>KY925611,<br>KY925636,<br>KY925736,<br>KY925871,<br>KY926039,<br>KY926287 | A/Torres/<br>LACENRS-1102/2016<br>KY925032,<br>KY925049,<br>KY925190,<br>KY925284,<br>KY925584,<br>KY926217,<br>KY926257,<br>KY926372 | A/Baltimore/<br>0107/2016<br>KY487084,<br>KY487183,<br>KY487439,<br>KY487451,<br>KY487472,<br>KY487556,<br>KY487562,<br>KY487722 | A/Taipei/<br>0046/2016<br>KY487170,<br>KY487436,<br>KY487504,<br>KY487531,<br>KY487560,<br>KY487635,<br>KY487734,<br>KY487740 | A/Baltimore/<br>0008/2016<br>KY487127,<br>KY487343,<br>KY487410,<br>KY487491,<br>KY487576,<br>KY487604,<br>KY487672,<br>KY487698 |
| 2 | A/Minnesota/<br>35/2017<br>CY243771 - 8 | A/Florida/<br>95/2015<br>KX004656 - 63 | A/Hawaii/<br>83/2015<br>KX004967 - 74 | A/Florida/<br>83/2015<br>KX005095,<br>KX005097 - 8,<br>KX005100,<br>KX005102,<br>KX005472,<br>KX005475,<br>KX005477 | A/Florida/<br>12/2016<br>KX406328 - 35 | A/Washington/<br>01/2017<br>CY218075 - 82 | A/North Carolina/<br>12/2016<br>KX407448 - 55 |
| 3 | A/Warsaw/<br>INS3_657/2011<br>CY176594 - 601 | A/Khon Kaen/<br>INS3_649/2012<br>CY176538 - 45 | A/Chicago/<br>YGA_04019/2012<br>CY171159 - 66 | A/Tennessee/<br>F2068/2011<br>CY167660 - 7 | A/Lima/<br>INS3_677/2012<br>CY176722 - 9 | A/Bronx/<br>INS3_673/2012<br>CY176690 - 7 | A/Georgia/<br>M5081/2012<br>CY147964,<br>CY147998,<br>CY148061,<br>CY148064,<br>CY148081,<br>CY148159,<br>CY148163,<br>CY148218 |
| 4 | A/Helsinki/<br>848/2013<br>KF560110 - 7 | A/Helsinki/<br>804/2013<br>KF560054 - 61 | A/Helsinki/<br>911/2013<br>KF560158 - 65 | A/Helsinki/<br>100/2013<br>KF559390 - 7 | A/Helsinki/<br>1199/2012<br>KF560278 - 85 | A/Helsinki/<br>205/2013<br>KF559438 - 45 | A/Helsinki/<br>737/2013<br>KF559990 - 7 |
| 5 | A/Parma/<br>644/2010<br>KU322148,<br>KU322190,<br>KU322233,<br>KU322276,<br>KU322319,<br>KU322362,<br>KU322405,<br>KU322448 | A/Parma/<br>7/2011<br>KU322178,<br>KU322220,<br>KU322263,<br>KU322306,<br>KU322349,<br>KU322392,<br>KU322435,<br>KU322478 | A/Parma/<br>48/2011<br>KU322176,<br>KU322218,<br>KU322261,<br>KU322304,<br>KU322347,<br>KU322390,<br>KU322433,<br>KU322476 | A/Parma/<br>126/2011<br>KU322173,<br>KU322215,<br>KU322258,<br>KU322301,<br>KU322344,<br>KU322387,<br>KU322430,<br>KU322473 | A/Singapore/<br>GP917/2011<br>CY124683 - 90 | A/Singapore/<br>GP887/2011<br>CY124663 - 70 | A/Singapore/<br>GP767/2011<br>CY124611 - 8 |
| 6 | A/Bangkok/<br>INS478/2010<br>CY096298 - 305 | A/Sydney/<br>DD3-37/2010<br>CY092638 - 45 | A/Texas/<br>JMS413/2010<br>CY061187 - 94 | A/Bangkok/<br>INS424/2010<br>CY071327 - 34 | A/Vienna/<br>INS368/2010<br>CY071055 - 62 | A/Berlin/<br>INS430/2010<br>CY071375 - 82 | A/Athens/<br>INS396/2010<br>CY071167 - 74 |
| 7 | A/NewYork/<br>A-WC-LVD-16-067/2016<br>CY259847 - 54 | A/Alaska/<br>263/2015<br>KU509765 - 7,<br>KU509771,<br>KX004187 - 8,<br>KX004696,<br>KX004702 | A/NewYork/<br>A-WC-LVD-16-040/2016<br>CY259655 - 62 | A/NewYork/<br>A-WC-LVD-16-018/2016<br>CY258913 - 20 | A/NewYork/<br>A-WC-LVD-16-016/2016<br>CY259463 - 70 | A/NewYork/<br>A-WC-LVD-16-053/2016<br>CY259751 - 8 | A/NewYork/<br>A-WC-LVD-16-014/2016<br>CY259447 - 54 |
| 8 | A/Mexico/<br>8017/2017<br>MF593572 - 9 | A/Mexico/<br>4436/2016<br>MF593514 - 21 | A/Mexico/<br>8517/2017<br>MF593580 - 7 | A/Mexico/<br>4687/2017<br>MF593551 - 8 | A/Mexico/<br>4435/2016<br>MF593506 - 13 | A/Mexico/<br>4440/2016<br>MF593522 - 9 | A/Mexico/<br>17517/2017<br>MF593617 - 24 |
| 9 | A/Thailand/<br>SN11783/2012<br>KP637595 - 602 | A/Milano/<br>21/2012<br>KU321908,<br>KU321939,<br>KU321971,<br>KU322003,<br>KU322033,<br>KU322065,<br>KU322097,<br>KU322129 | A/Taiwan/<br>1829/2011<br>JQ693691,<br>JQ693701,<br>JQ693711,<br>JQ693721,<br>JQ693731,<br>JQ693741,<br>JQ693751,<br>JQ693761 | A/Indiana/<br>04/2011<br>KC882315,<br>KC882319 - 21,<br>KC882343 - 5,<br>KC882371 | A/Michigan/<br>03/2011<br>KC882264,<br>KC882275 - 80,<br>KC882296 | A/Pennsylvania/<br>07/2011<br>KC881821 - 4,<br>KC882000,<br>KC882022 - 3,<br>KC882124 | A/Budapest/<br>WRAIR3789T/2011<br>CY097977 - 84 |

**Supplemental Table S5.**
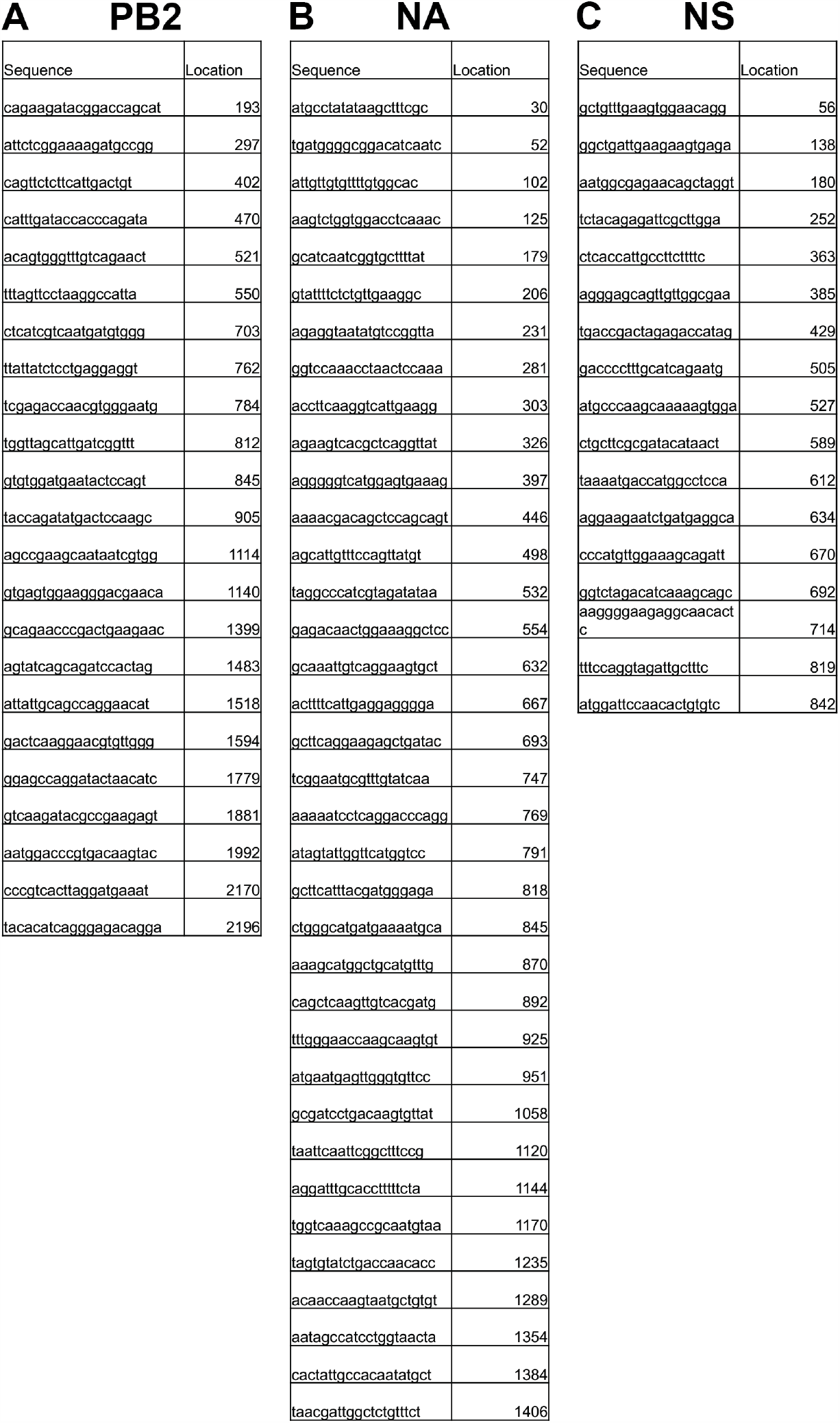
FISH probe sequences. Custom oligonucleotide probes targeting PB2 **(A)**, NA **(B)**, and NS **(C)** vRNA were designed from A/Panama/2007/1999 (H3N2) virus sequences using the Stellaris probe designer (BioSearch Technologies). Oligos exhibiting significant complementarity against other vRNA segments and/or positive-strand complemen tarity were excluded.

## Acknowledgments

All confocal microscopy imaging was performed at the Center for Biologic Imaging at the University of Pittsburgh. JEJ is supported by a T32 (T32 AI049820) and the Catalyst Award (University of Pittsburgh Center for Evolutionary Biology and Medicine). This work is funded by the National Institutes of Health NIAID (R01 AI139063). We thank members of the Lakdawala and Wright labs for technical support and constructive feedback on this manuscript.

